# A Primase-Induced Conformational Switch Controls the Stability of the Bacterial Replisome

**DOI:** 10.1101/469312

**Authors:** Enrico Monachino, Slobodan Jergic, Jacob S. Lewis, Zhi-Qiang Xu, Allen T.Y. Lo, Valerie L. O’Shea, James M. Berger, Nicholas E. Dixon, Antoine M. van Oijen

## Abstract

Recent studies of bacterial DNA replication have led to a picture of the replisome as an entity that freely exchanges DNA polymerases and displays intermittent coupling between the helicase and polymerase(s). Challenging the textbook model of the polymerase holoenzyme acting as a stable complex coordinating the replisome, these observations suggest a role of the helicase as the central organizing hub. We show here that the molecular origin of this newly-found plasticity lies in the >400-fold increase in strength of the interaction between the polymerase holoenzyme and the replicative helicase upon association of the primase with the replisome. By combining *in vitro* ensemble-averaged and single-molecule assays, we demonstrate that this conformational switch operates during replication and promotes recruitment of multiple holoenzymes at the fork. Our observations provide a molecular mechanism for polymerase exchange and offer a revised model for the replication reaction that emphasizes its stochasticity.

## INTRODUCTION

The *Escherichia coli* replisome is composed of at least 12 individual proteins that work together to coordinate leading- and lagging-strand synthesis during the copying of chromosomal DNA (reviewed in Lewis et al., 2016) (Figure 1A). Following unwinding of the parental double-stranded (ds) DNA by the DnaB helicase, the DNA polymerase III holoenzyme (Pol III HE) synthesizes DNA on the two daughter strands. The single-stranded (ss) leading strand is displaced by the DnaB helicase and duplicated continuously, while the lagging strand is extruded through the DnaB central channel, coated by ssDNA-binding protein (SSB) and converted to dsDNA discontinuously during the production of 1–2 knt Okazaki fragments (OFs) (Kornberg and Baker, 1991). The distinct asymmetry in the mechanism of synthesis of the two strands finds its origin in their opposite polarity, requiring that lagging-strand synthesis take place in a direction opposite to that of synthesis of the leading strand. This situation is proposed to result in the formation of a lagging-strand loop (Sinha et al., 1980).

**Figure 1.**
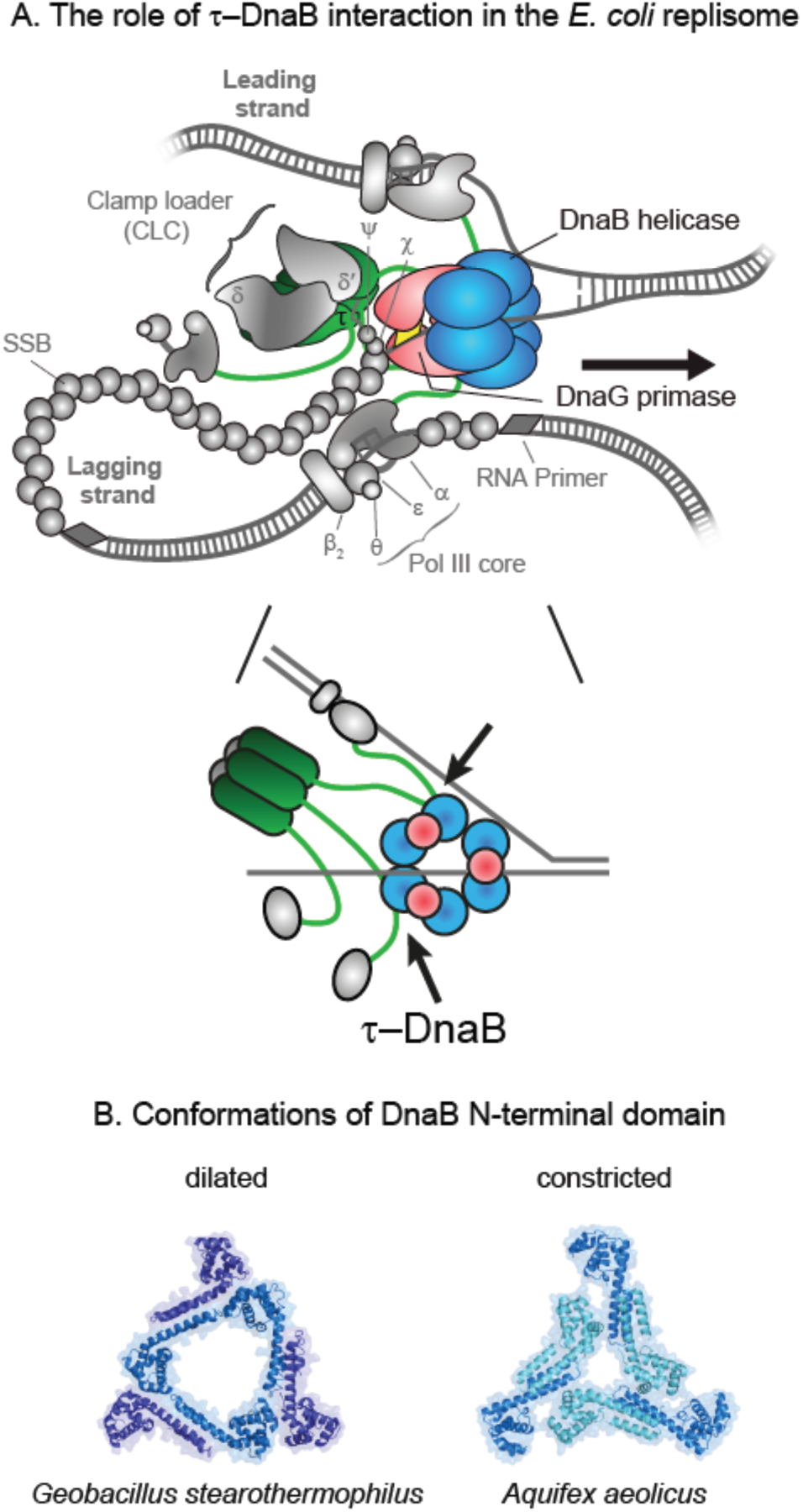
DnaB and Its Binding Partners in the *E. coli* Replisome. (A) Schematic representation of the *E. coli* replisome (top panel). DnaB, DnaG, and τ are shown in color, with the other components of the elongating replisome (αεθ Pol III cores, β_2_ sliding clamps, SSB, and CLC subcomplexes δδ’ and *χ*ψ) presented in shades of gray. The schematic in the bottom panel provides a flattened representation of the key interactions between DnaB, DnaG, and τ. (B) The two conformations of DnaB helicase N-terminal domains – dilated (left panel; PDB: 2R6A) and constricted (right panel; PDB: 4NMN).

Each OF is initiated at a short RNA primer deposited by the DnaG primase for utilization by Pol III HE. Primase requires interaction with the helicase to stimulate its RNA polymerase activity (Johnson et al., 2000). The DnaB–DnaG contact is established through the interaction between the C-terminal domain of primase, termed DnaGC (Tougu and Marians, 1996; Oakley et al., 2005), and the N-terminal domains of the helicase (Bailey et al., 2007). In *E. coli*, this interaction is weak and transient, with *K*_D_ in the low μM range and fast on/off kinetics (Oakley et al., 2005), whereas in *Geobacillus stearothermophilus*, DnaB and DnaG form a stable complex that can be isolated by gel filtration (Bird et al., 2000) and crystallized (Bailey et al., 2007).

In bacteria, DnaB-family helicases form homo-hexamers with a distinct two-layered ring structure. The C-terminal RecA-like AAA ATPase domains that power DNA unwinding have pseudo-six-fold symmetry. In contrast, the N-terminal domains typically display three-fold (C3) symmetry, established from a trimer-of-dimers that encircle a wide, open (dilated) central channel that can accommodate dsDNA (Figure 1B, left panel) (Bujalowski et al., 1994; San Martin et al., 1995; Yu et al., 1996; Bailey et al., 2007; Strycharska et al., 2013). However, a crystal structure of *Aquifex aeolicus* DnaB in the presence of nucleotides showed a different C3 arrangement in the N-terminal collar whereby the central pore is narrow (constricted) and only large enough to accommodate ssDNA. Associated electron microscopy studies showed that *E. coli* DnaB can adopt this state as well (Figure 1B, right panel) (Strycharska et al., 2013). These two strikingly different structures suggest that a relatively low energy barrier exists between the dilated and constricted states and that the helicase may transition between them as it translocates along ssDNA.

The Pol III HE, the complex responsible for the extension of deposited RNA primers and most of chromosomal DNA synthesis, interacts with the replicative DnaB helicase (Kim et al., 1996; Dallmann et al., 2000; Gao and McHenry, 2001a). The Pol III HE is composed of three functionally distinct subassemblies that can be isolated separately from individual subunits: the Pol III core (or just Pol III), the sliding clamp, and the clamp loader complex (CLC) (Kelman and O’Donnell, 1995). The catalytic Pol III cores are heterotrimers of α, ε and θ subunits responsible for DNA synthesis and proofreading (Scheuermann and Echols, 1984; Maki and Kornberg, 1985; Studwell-Vaughan and O’Donnell, 1993; Taft-Benz and Schaaper, 2004). The sliding clamp is a toroid-shaped β_2_ homodimer (Kong et al., 1992). Once loaded on the DNA by the ATP-dependent activity of the CLC, the clamp stabilizes Pol III on the DNA template and improves processivity through a pair of weak interactions with the α and ε subunits (Naktinis et al., 1996; Dohrmann and McHenry, 2005; Jergic et al., 2013; Fernandez-Leiro et al., 2015). The CLC contains seven proteins and has the composition *χ*ψτ*nγ*_(3–*n*)_δδ’; *n* = 0–3, where physiologically relevant assemblies are thought to have *n* = 2 or 3 τ subunits (Reyes-Lamothe et al., 2010; Lewis et al., 2016). The unique δ and δ’ subunits interact with the three copies of the *dnaX* gene product oligomer, τ and/or *γ* (Kodaira et al., 1983; Mullin et al., 1983) to assemble into a stable ATPase-proficient semi-circular pentamer (Jeruzalmi et al., 2001; Bullard et al., 2002; Simonetta et al., 2009). Whereas τ is the full-length product of *dnaX*, *γ* is a C-terminally truncated version produced as a result of a programmed ribosomal frame-shift during mRNA translation (Tsuchihashi and Kornberg, 1990; Flower and McHenry, 1990; Blinkowa and Walker, 1990). The accessory subunits *χ* and ψ form a strong heterodimeric complex that interacts with all three τ/*γ* subunits of the pentamer via the flexible N-terminal residues in ψ (Gulbis et al., 2004; Simonetta et al., 2009) to assemble the full CLC.

The τ subunit provides the physical connectivity between the polymerase and helicase activities. It has a five-domain structure (Gao and McHenry, 2001b), with the N-terminal domains I–III being identical to *γ* and responsible for oligomerization and ATPase-dependent clamp loading activity. The C-terminal fragment that distinguishes τ from *γ* is termed τ_c24_. It contains domain IV, which transiently engages DnaB as a monomer (Gao and McHenry, 2001a), and domain V, which provides a strong, slowly-dissociable interaction with α (Gao and McHenry, 2001b; Jergic et al., 2007). Consequently, the τ subunit of the CLC plays a key linking role in the replisome: it ensures cohesion of the Pol III–CLC particle (αεθ)*_n_*–χψτ_*n*_*γ*_(3–*n*)_δδ’ (*n* = 2–3), termed Pol III^∗^, and links the complex to the replicative helicase (Figure 1A).

Recent advances in the field have challenged the deterministic view of the replisome as a perfectly orchestrated machine, whereby a single Pol III^∗^, stably bound to the replication fork, replicates DNA in a strictly ordered sequence of events (van Oijen and Dixon, 2015; Monachino et al., 2017; Graham et al., 2017). Instead, frequent turnover of the Pol III^∗^ in the replisome has been observed (Yuan et al., 2016; Beattie et al., 2017; Lewis et al., 2017), suggesting that the helicase acts as the central organizing structure of the replisome as opposed to the polymerase. An explanation for this surprising level of plasticity can be found in the network of weak interactions that enable polymerases from solution to eventually replace those at the fork (Geertsema and van Oijen, 2013; Lewis et al., 2017). However, this explanation seems at odds with the strong and stable interaction between multimeric τ (Pritchard et al., 2000; Park et al., 2010) and DnaB (Kim et al., 1996; Gao and McHenry, 2001a).

We present here an unexpected conformational switch in the DnaB helicase that is triggered upon binding of DnaG, that increases the strength of the DnaB–CLC interaction by more than two orders of magnitude. We find that in solution, DnaB is almost exclusively in the constricted state, as observed in the crystal structure of *Aquifex aeolicus* DnaB. However, during replication, it transits between the dilated and constricted states in a manner that is controlled by
primase interaction. Nevertheless, DnaB remains an active helicase as it samples both states. Finally, we use single-molecule visualization of Pol III during replication to show that the primase concentration modulates both the kinetics of polymerase exchange in the replisome and the steady-state number of polymerases associated with the replication fork.

Taken together, our observations establish a model whereby the interaction between the primase and helicase acts as a switch to control the organization and dynamics of the replisome. This realization has important ramifications for understanding of coordination of leading- and lagging-strand synthesis, the coupling between polymerase and helicase, and the timing of Okazaki-fragment synthesis.

## RESULTS

### χψ Serves as a Molecular Anchor for the Clamp-Loader Complex

Due to its weak nature and complex stoichiometry, the interaction between multiple τ subunits in the CLC and DnaB is poorly understood. Previous studies employed surface plasmon resonance (SPR) to identify the region within τ that is responsible for binding to DnaB (Gao and McHenry, 2001a), but the use of monomeric τ fragments immobilized on the surface makes it challenging to interpret these results in the context of multiple τ subunits within the CLC interacting with DnaB simultaneously. To study these interactions in a context that closely represents the physiologically relevant system, we immobilized the entire CLC onto a streptavidin (SA)-coated chip surface through an N-terminally biotinylated *χ* subunit (^bio^*χ*), and used this surface-immobilized CLC as a platform to measure the interaction with the helicase.

The stability of the CLC on the SA chip was first assessed by monitoring the dissociation of *in situ* assembled τ_3_CLC from immobilized ^bio^*χ*ψ and associated τ_3_δδ’ (Figures S1A–C). In a high ionic strength buffer (200 mM NaCl), dissociation was moderately slow (a dissociation half-life *t*_1/2_ of ~50 min) and was unaffected by the presence of 1 mM ADP (Figure S1C). In addition, the dissociation of ψ from immobilized ^bio^*χ* was slow (Figure S1B) and re-injection of same concentration of τ_3_δδ’ (100 nM) after 2 days led to 50% recovery of the original signal (not shown), indicating that the population of ψ still bound to ^bio^*χ* was only halved, suggesting a *t*_1/2_ for the *χ*–ψ interaction of ~2 days. A short injection of 1 M MgCl_2_ resulted in loss of ~25% of mass from the surface (Figure S1D) and initiated a faster dissociation phase (*t*_1/2_ ~10 min), which could be slowed down to the original level if both δ and δ’ (but not δ’ alone) were injected over the surface (*t*_1/2_ ~ 45 min; Figure S1 E,F). We did not inject δ alone because it interacts weakly with τ/*γ* in the absence of δ’ (Onrust et al., 1995). The data indicated that the treatment with MgCl_2_ could not regenerate *χ*ψ on the surface, but rather led to separation of δδ’, followed by ~five-fold faster dissociation of τ_3_ from ψ (Simonetta et al., 2009). Considering that δδ’ does not contact ψ directly (Glover and McHenry, 1998; Simonetta et al., 2009), the change in dissociation rate revealed that δδ’ contributes to the proper conformation of τ_3_ in the CLC for the interaction with ψ. Taken together, our data are consistent with the observation of wholesale dissociation of the entire τ_3_δδ’ from *χ*ψ in buffer with 200 mM NaCl, thus pointing to the ψ–(τ/*γ*)_3_ interaction being the weakest link in ^bio^CLC.

Determination of the stability of the CLC on the SPR chip surface was necessary prior to further binding studies. Since we determined its stability is directly related to the slow dissociation of τ_3_ from ψ, and considering that ^bio^*χ*ψ could not be efficiently regenerated, we set out to investigate the rates of *χ*ψ-*γ*_3_δδ’/τ_2_*γ*_1_δδ’ interactions using single-shot kinetics on a multiplexed ProteOn SPR system. Previous SPR measurements have indicated that the *K*_D_ for the *χ*ψ–τ/*γ* interaction is ~2 nM at relatively low ionic strength (100 mM potassium glutamate, K-Glu) (Olson et al., 1995). Our analysis indicated that the ψ–(τ/*γ*)_3_ interaction should be stronger if (τ/*γ*)_3_ is part of a pentamer with δδ’. We measured the binding kinetic parameters for *χ*ψ-*γ*_3_δδ’ and *χ*ψ-τ_2_*γ*_1_δδ’ interactions by injecting various concentrations of CLC cores over immobilized ^bio^*χ*ψ in a buffer containing 200 mM NaCl. Irrespective of the CLC core used, we measured a dissociation *t*_1/2_ of ~50 min (Table 1; Figure S2A,B, top panels), and dissociation constants of ~1 and ~2 nM for *γ*_3_δδ’ and τ_2_*γ*_1_δδ’ from ^bio^*χ*ψ, respectively.

**Table 1:** Binding Parameters for the ^bio^*χ*ψ–*γ*_3_δδ’ and ^bio^*χ*ψ–τ_2_*γ*_1_δδ’ Interactions, With or Without ATP. Equilibrium constant (*K*_D_), and association (*k*_a_) and dissociation (*K*_d_) rate constants, including the calculated dissociation half-life (*t*_1/2_), were determined by simultaneous fitting of sensorgrams in Figure S2 to 1:1 (Langmuir) binding model. The errors are standard errors of the fits. See also Figures S1 and S2.

| Interaction | $K_D$ (nM) | $k_a$ ($M^{-1}s^{-1}$) | $k_d$ ($s^{-1}$) | $t_{1/2}$ (min) |
| --- | --- | --- | --- | --- |
| $^{bio}\chi\psi-\gamma_3\delta\delta'$ (ATP<sup>-</sup>) | $1.1 \pm 0.0$ | $(2.06 \pm 0.00) \cdot 10^5$ | $(2.19 \pm 0.00) \cdot 10^{-4}$ | 53 |
| $^{bio}\chi\psi-\tau_2\gamma_1\delta\delta'$ (ATP<sup>-</sup>) | $2.2 \pm 0.0$ | $(9.59 \pm 0.01) \cdot 10^4$ | $(2.10 \pm 0.00) \cdot 10^{-4}$ | 55 |
| $^{bio}\chi\psi-\gamma_3\delta\delta'$ (ATP<sup>+</sup>) | $0.3 \pm 0.0$ | $(2.67 \pm 0.00) \cdot 10^5$ | $(8.10 \pm 0.01) \cdot 10^{-5}$ | 143 |
| $^{bio}\chi\psi-\tau_2\gamma_1\delta\delta'$ (ATP<sup>+</sup>) | $0.3 \pm 0.0$ | $(1.71 \pm 0.00) \cdot 10^5$ | $(5.40 \pm 0.01) \cdot 10^{-5}$ | 214 |

Searching for a further increase of CLC stability on the chip surface, we investigated the effect of nucleotides. ADP had no effect on dissociation of ^bio^CLC (Figure S1C); however, as the ψ subunit is known to stabilize the ATP-induced conformational state of the CLC (Anderson et al., 2007; Simonetta et al., 2009), ATP could be expected to stabilize the construct and in particular the ψ–(τ/*γ*)_3_ link according to the principle of microscopic reversibility. Indeed, we found this to be true: 1 mM ATP increased the affinity of *γ*_3_δδ’/τ_2_*γ*_1_δδ’ for ^bio^*χ*ψ by 3.5- and 7-fold (to ~0.3 nM) and increased the dissociation *t*_1/2_ 3- and 4-fold, respectively (Table 1; Figure S2A,B). By reducing the ionic strength to 50 mM NaCl and by chromatographically isolating the entire ^bio^CLCs prior to immobilization (Figure S3A), we further improved the stability of the CLC on the chip surface. A dissociation *t*_1/2_ of >10 h (Figure S3B) provided us with an ideal platform to study interactions between the immobilized CLC and DnaB.

### Clamp Loader-Helicase Affinity Increases >400-Fold upon DnaGC Binding

We next used SPR to test the strength of interaction between wild-type DnaB^wt^ and surface-immobilized τ_3_CLC (Figure 2A,B). Sensorgrams recorded at a range of concentrations of DnaB^wt^ injected over ^bio^*χ*ψτ_3_δδ’ in a buffer containing 1 mM ADP revealed unexpectedly fast kinetics, with fast on- and off-rates (Figure 2B). Using ATP instead of ADP did not affect responses at equilibrium (not shown) but was found to increase nonspecific surface interactions at high DnaB concentrations, which became a critical obstacle to measurements since CLCs on the chip surface could not be regenerated. Responses measured at equilibrium were fit to a 1:1 steady-state affinity (SSA) model (Equation 1, Methods) to yield a *K*_D_ value of 1.3 ± 0.2 μM. Similar measurements revealed an almost identical strength and similarly fast kinetics of the ^bio^*χ*ψτ_1_*γ*_2_δδ’–DnaB interaction (*KD* = 4.1 ± 0.3 μM; Figure S3C). Thus, unlike the expectation based on previously published results (Kim et al., 1996; Gao and McHenry, 2001a), our data show that the interaction between CLC and DnaB^wt^ is weak and transient and does not depend on the number of τ subunits in the CLC.

**Figure 2.**
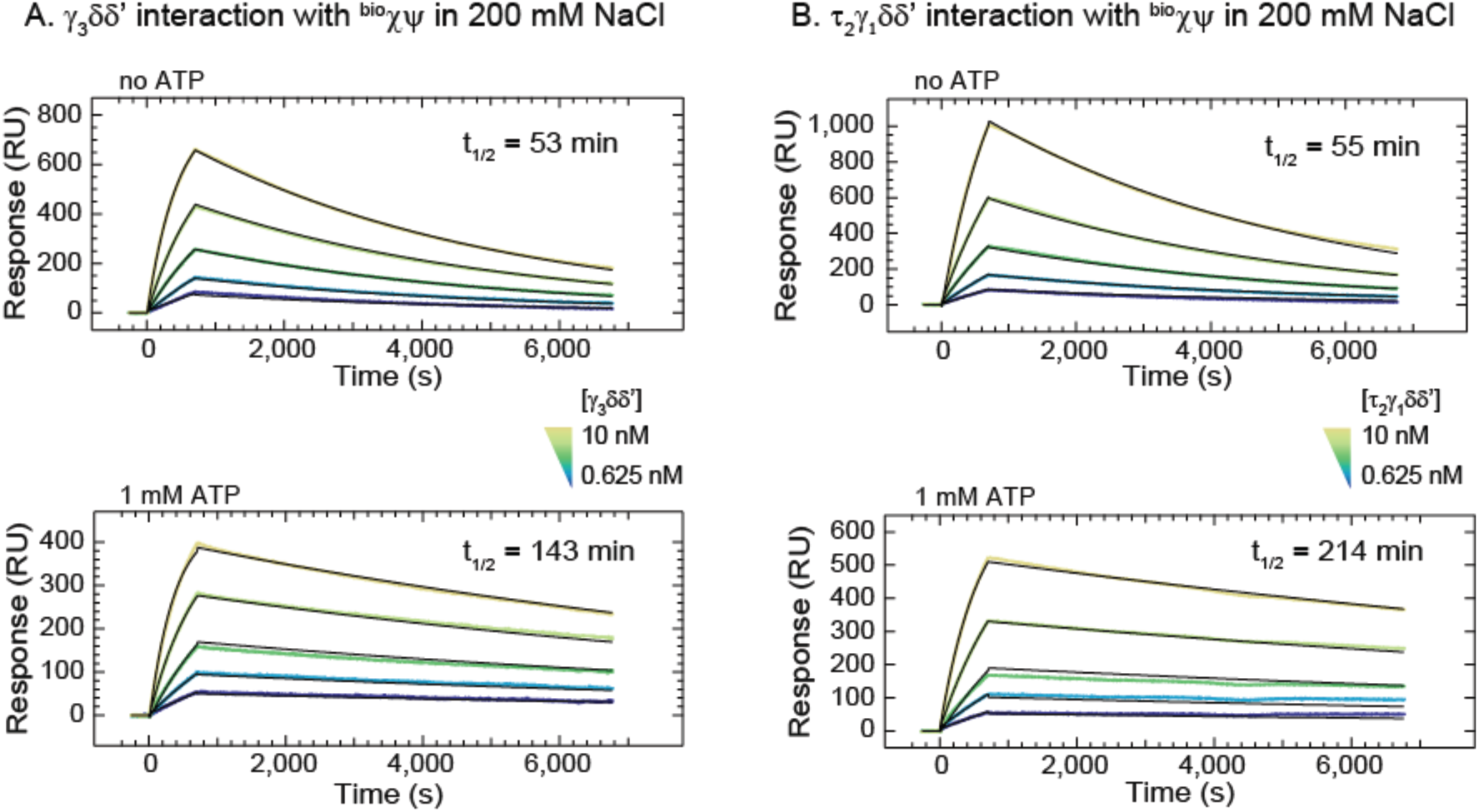
The Dilated Form of DnaB Interacts Tightly with ^bio^*χ*ψτ_3_δδ’. (A) Cartoon representation of the association and dissociation phases of an SPR experiment used to measure the binding of DnaB versions in solution to immobilized τ_3_CLC. (B) and (C) SPR sensorgrams show association (for 30 s) and dissociation phases of DnaB^wt^ from ^bio^*χ*ψτ_3_δδ’ (B) and DnaB^constr^ from ^bio^*χ*ψτ_3_δδ’ (C) for interactions obtained over a 0.0625–8 μM range of serially diluted DnaB^wt^ (B) or DnaB^constr^ (C), including a zero concentration control. Responses at equilibrium, determined by averaging values in the gray bar region, were fit (inset, red curve) using a 1:1 steady state affinity (SSA) model to derive dissociation constant *K*_D_ and response at saturation, *R*_max_: *K*_D_(^bio^*χ*ψτ_3_δδ’–DnaB^wt^) =1.3 ± 0.2 μM and *R*_max_ = 440 ± 20 RU (B), and *K*_D_(^bio^*χ*ψτ_3_δδ’–DnaB^constr^) = 3.3 ± 0.2 μM and *R*_max_ = 460 ± 20 RU (C). Errors are standard errors of the fit. (D) 250 nM DnaB^dilated^ is injected for 400 s and its slow dissociation monitored over 2,000 s. (E) Cartoon representation of bulk replication assays with rolling-circle substrates in the absence or presence of DnaG primase, as shown in (F). The presence of DnaG enables also the lagging-strand synthesis. (F) Alkaline agarose gel showing the resolved leading strand (all lanes) and lagging strand (DnaG^+^ lanes) DNA products generated by replisomes either in the absence of primase (odd lanes) or its presence (even lanes), using DnaB^dilated^ (lanes 1–2), DnaB^wt^ (lanes 3–4), and DnaB^constr^ (lanes 5–6) helicases. See also Figure S3.

Recent studies proposed that the τ subunit is able to differentiate between dilated and constricted DnaB states (Strycharska et al., 2013). This conclusion was based on DNA-unwinding assays performed with two mutant versions of the helicase that were stabilized in either the dilated (DnaB^dilated^) or constricted (DnaB^constr^) states (Figure 1B). We tested both mutants for their interaction with surface-immobilized τ_3_CLC using SPR. Whereas DnaB^constr^ exhibited binding kinetics and strengths similar to that of DnaB^wt^ (*K*_D_ = 3.3 ± 0.3 μM; Figure 2C), an injection of 250 nM DnaB^dilated^ resulted in a markedly slower dissociation from ^bio^*χ*ψτ_3_δδ’ (Figure 2D). The interaction was stabilized to a level that prevented us from reliably quantifying its strength, with the dissociation of DnaB^dilated^ from ^bio^*χ*ψ*γ*τ_3_δδ’ now competing with the dissociation of τ_3_δδ’ from ^bio^*χ*ψ. Nevertheless, the similarity of the kinetics of DnaB^wt^ and DnaB^constr^ interacting with τ_3_CLC and the stark difference between those of DnaB^wt^ and DnaB^dilated^ suggest that *E. coli* DnaB^wt^ is almost entirely in the constricted state in solution.

In a co-crystal structure of the *Geobacillus stearothermophilus* DnaB_6_•(DnaGC)_3_ complex, the three C-terminal domains of primase were each seen to bind to two adjacent subunits of the dilated hexameric collar domain in DnaB (Figure S4A). However, in the constricted conformation of *Aquifex aeolicus* DnaB (Figure 1B, right panel), one of the two asymmetric DnaGC contact points in DnaB is buried and unavailable for binding to DnaG (Strycharska et al., 2013). In agreement with the structural considerations, it was further reported that DnaB^dilated^ is able to interact with DnaG and support priming activity whereas DnaB^constr^ could not sustain priming at all. We compared the activities of wild-type DnaB and the two mutants in a bulk leading- and lagging-strand replication assay (Figure 2E, DnaG^+^ path) and found that, unlike DnaB^wt^ and DnaB^dilated^, DnaB^constr^ was indeed incapable of sustaining OF synthesis upon addition of DnaG (Figure 2F; lane 6 *cf.* lanes 2 and 4). By contrast, all three helicases, including DnaB^constr^ were proficient in leading-strand synthesis (Figure 2E, DnaG^−^ path, and Figure 2F, lanes 1, 3, and 5). These findings demonstrate that that the dilated and constricted states of DnaB both support helicase activity. Considering that DnaB^wt^ is largely in the constricted state in solution, the observation that DnaB^wt^ supports the production of OFs also indicates that the helicase is able to explore dilated-like states during replication, due either to primase or ssDNA binding, or both.

To test the possibility that DnaG binding to DnaB is sufficient to trigger the conformational transition in the helicase, we produced DnaGC (Loscha et al., 2004), a C-terminal domain of primase that has all the determinants for DnaB binding but lacks the two N-terminal domains responsible for recognizing priming sites on DNA and for RNA synthesis. Both DnaG and DnaGC interact similarly weakly with DnaB^wt^ with *K*_D_ values of 2.8 and 4.9 μM in 150 mM NaCl, respectively (Oakley et al., 2005). Injection of 0.5 μM DnaB^wt^ together with 5 μM DnaGC and 1 mM ATP (50 mM NaCl) over immobilised τ_3_CLC (Figure S4B) resulted in a much higher response compared to the injection of DnaB^wt^ alone (Figure S4B). This stronger binding was not ATP specific, since the use of ADP resulted in the similar response (Figure S4C). Injection of 5 μM DnaGC alone did not yield a detectable response, showing that the signal is not caused by direct binding of DnaGC to τ_3_CLC (Figure S4B). Critically, fast-off kinetics were detected when DnaGC was omitted during the dissociation of DnaB from the CLC, as if the DnaGC was not present during the association phase at all (Figures S4B *cf.* 2B). Considering that DnaGC appears to dramatically increase the affinity of DnaB for CLC, the fast dissociation detected appeared to violate the thermodynamic principle that each ligand (CLC or DnaGC) must increase the affinity of protein (DnaB) for the other. To further investigate this behavior, we measured the binding of DnaB^wt^ to immobilised τ_3_CLC by fixing the solution DnaB^wt^ concentration to 100 nM while titrating DnaGC over a range of 0.5–8 μM (Figure S4D). Despite the presence of three binding sites on DnaB for DnaGC, the data were readily fit by a simple SSA model of one-to-one binding, as if only one of the three sites on DnaB had been bound by DnaGC in the measured concentration range. The measured *K*_D_ of 1.74 ± 0.09 μM and the fast kinetics were similar to the previously reported binding of DnaB^wt^ to individual immobilized DnaG subunits (Oakley et al., 2005). These observations can be reconciled by a model that describes a cooperative transition in DnaB, with the weak binding of the first DnaGC initiating a conformational transition in the DnaB hexamer from a constricted to a dilated state that has at least a 10–100-fold higher affinity for a second (and possibly third) DnaGC.

The strength of the interaction between τ_3_CLC and the DnaB^wt^•DnaGC complex can be now determined accurately by measuring responses at equilibrium for various concentrations of DnaB^wt^ in the presence of 5 μM DnaGC and fitting against the calculated concentration of DnaB^wt^•DnaGC (see Equation 2, Methods) using an SSA model (*K*_D_ = 2.6 ± 0.3 nM; Figure 3A). Likewise, binding of DnaB^wt^ to τ_1_CLC at 5 μM DnaGC was similarly strong (*K*_D_ = 10 ± 1 nM; Figure S4E), again arguing that a single τ subunit within the τ_3_CLC is responsible for helicase binding. Nevertheless, the increase in the strength of DnaB^wt^–CLC interaction by up to 400-fold in the presence of DnaGC indicates that the binding of the primase triggers a conformational switch in DnaB from the constricted to a dilated state, stabilizing its binding to the CLC. As expected, injection of DnaB^constr^ in the presence of DnaGC resulted in a much weaker response, consistent with the incompatibility of this conformation to stably interact with primase (Figure S4F).

**Figure 3.**
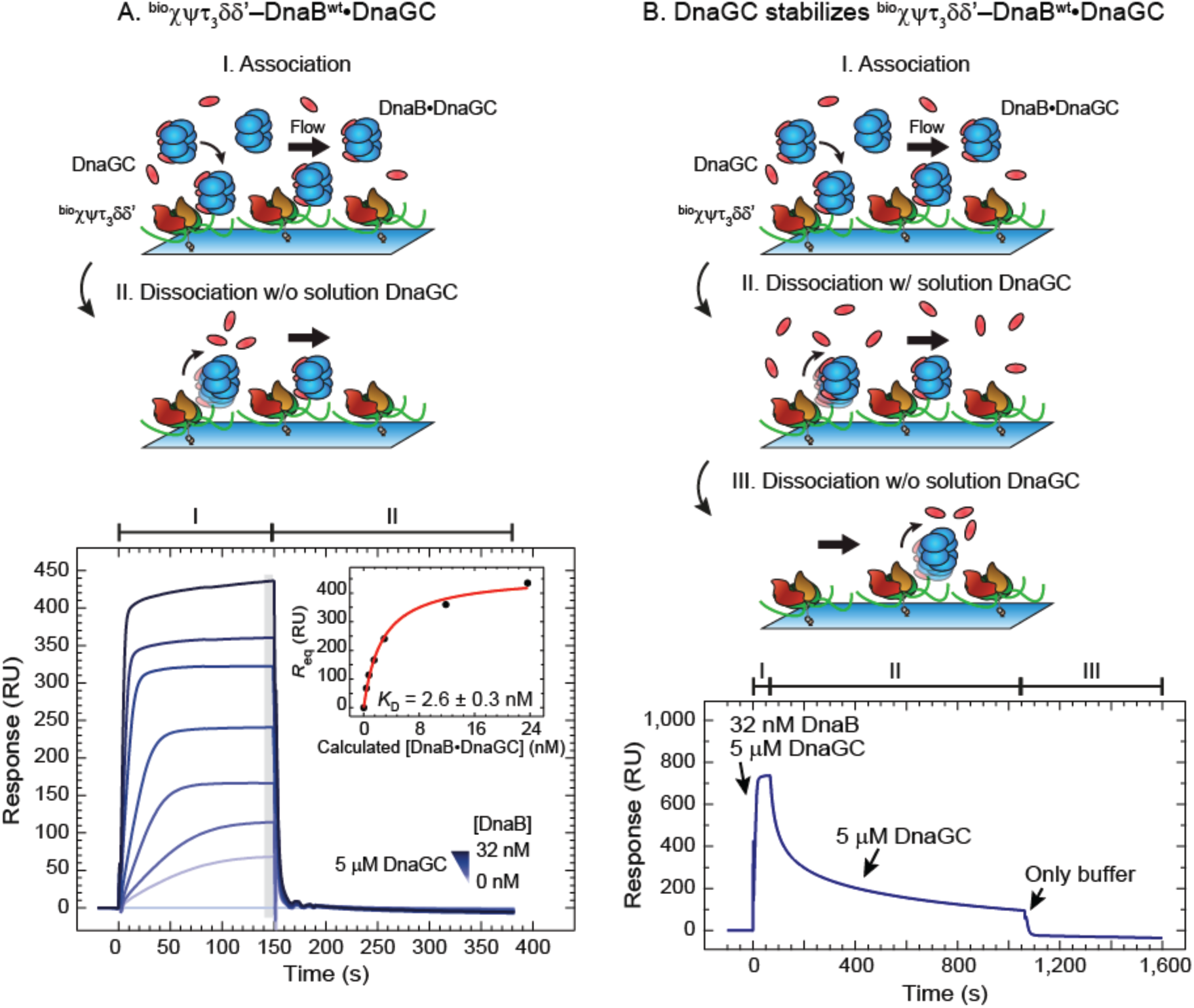
Association of DnaGC with DnaB Strengthens the ^bio^*χ*ψτ_3_δδ’–DnaB Interaction 500-Fold. (A) *Top panel*: Cartoon representation of the association and dissociation phases of an SPR experiment used to measure the binding of solution DnaB^wt^ to immobilized τ_3_CLC in the presence of DnaGC during association only. *Bottom panel:* SPR sensorgrams showing association (for 150 s) and dissociation of DnaB•DnaGC to and from ^bio^*χ*ψτ_3_δδ’ obtained over a 0.5–32 nM range of serially-diluted DnaB^wt^ (including 0 nM control) together with 5 μM DnaGC. The responses at equilibrium *R*_eq_, determined by averaging values in the gray bar regions of the sensorgrams, were fit (inset, red curve) against the calculated DnaB^wt^•DnaGC concentrations (0–23.7 nM, see Methods) using an SSA model to obtain a *K*_D_ value of 2.6 ± 0.3 nM and an *R*_max_ value of 460 ± 10 RU. Errors are standard errors of the fit. (B) *Top panel:* Cartoon representation of DnaB^wt^ association (with DnaGC, step I) and dissociation (with, step II, and without DnaGC, step III, respectively) during an SPR experiment. *Bottom panel*: DnaGC slows the dissociation of DnaB^wt^ from ^bio^*χ*ψτ_3_δδ’. SPR sensorgram shows the association of DnaB^wt^•DnaGC during a 60 s injection of 32 nM DnaB^wt^ in the presence of 5 μM DnaGC (step I), followed by a 1,000 s dissociation phase in the presence of 5 μM DnaGC (step II) and second, rapid dissociation phase without DnaGC (step III). See also Figure S4.

Interestingly, the presence of DnaGC (5 μM) in the dissociation phase slows down τ_3_CLC•DnaB dissociation to a lifetime of several 100s of seconds (Figure 3B). These observations indicate that the DnaB conformation is regulated by primase–helicase interaction, thereby controlling the affinity of DnaB to the CLC. On the other hand, the CLC cannot lock DnaB in its dilated state. We thus identified two functional forms of the helicase–clamp loader interaction: one with a weak affinity with the helicase in the constricted-like state and the other with a strong affinity that depends exclusively on cooperative primase–helicase interactions.

### The Strong Helicase–Clamp Loader Interaction Stimulates the Activity of a Destabilized Replisome

The τ subunit of the CLC has a central role in the replisome, connecting the polymerase core with the replicative helicase. Hence, a primase-induced >400-fold increase in DnaB–CLC affinity and the strongly altered kinetics of the formation of this complex could be expected to significantly affect the organization and dynamics of the replisome. To demonstrate the importance of the strong helicase–clamp loader interaction in a functional context, we first turned to a rolling-circle leading-strand bulk replication assay (Figure 2E, DnaG^−^ path). In this well-established assay (Mok and Marians, 1987), the β_2_–Pol NI^lead^–CLC–DnaB connectivity stimulates the simultaneous unwinding of dsDNA by DnaB and primer extension DNA synthesis by a Pol NI^lead^ core bound to the leading strand. We modified the assay such that the stability of Pol III^∗^ on DNA is compromised by leaving out the β_2_ processivity factor in the reaction (Figure 4A). This condition artificially elevates the importance of the remaining DnaB–τ_3_δδ’ link and allows us to visualize its functional dependence on the DnaGC concentration, [DnaGC]. Because of the expected inefficiency of the reaction, we first performed a time-course assay at constant DnaGC (2 μM) (Figure 4B, lanes 2–5). We found that the reaction still progresses and that the products are best observable at 80 min, with progressively longer products synthesized and more DNA templates consumed in time. Moreover, in the absence of DnaGC, replication was significantly less efficient and equivalent to the level in the DnaGC-dependent reaction at early time points (Figure 4B, lane 6 *cf.* lanes 2 and 3). The narrow distribution of product sizes points to the distributive nature of the DNA-synthesis process, not surprisingly considering the absence of β_2_. Performing the reaction at progressively increasing [DnaGC] confirms a dependence of the synthesis efficiency on DnaGC (Figure 4C, lanes 1–5). Considering that (i) DnaGC has no enzymatic activity, (ii) leading-strand replication does not proceed in the absence of the physical coupling between the helicase and Pol III^∗^ (Kornberg and Baker, 1991; Kim et al., 1996), (iii) helicase-independent Pol III strand displacement synthesis cannot occur in the absence of β_2_ (Yuan and McHenry, 2009; Jergic et al., 2013), (iv) the efficiency of replication increases in the concentration range of DnaGC that is relevant for its interaction with DnaB (Figure S4D) and stabilization of the DnaB–CLC interaction (Figure 3), (v) helicase loading, presumably occurring by sliding onto the free 5’-end of the lagging strand, was unaffected by increasing [DnaGC] (as observed by a constant template utilization as a function of [DnaGC]), and (vi) DnaB is reported to be stable on ssDNA for very long periods (~60 min; Pomerantz and O’Donnell, 2010), we conclude that the progressively higher efficiencies in DNA synthesis are due to a DnaGC-induced strengthening of the Pol III^∗^–DnaB interaction.

**Figure 4.**
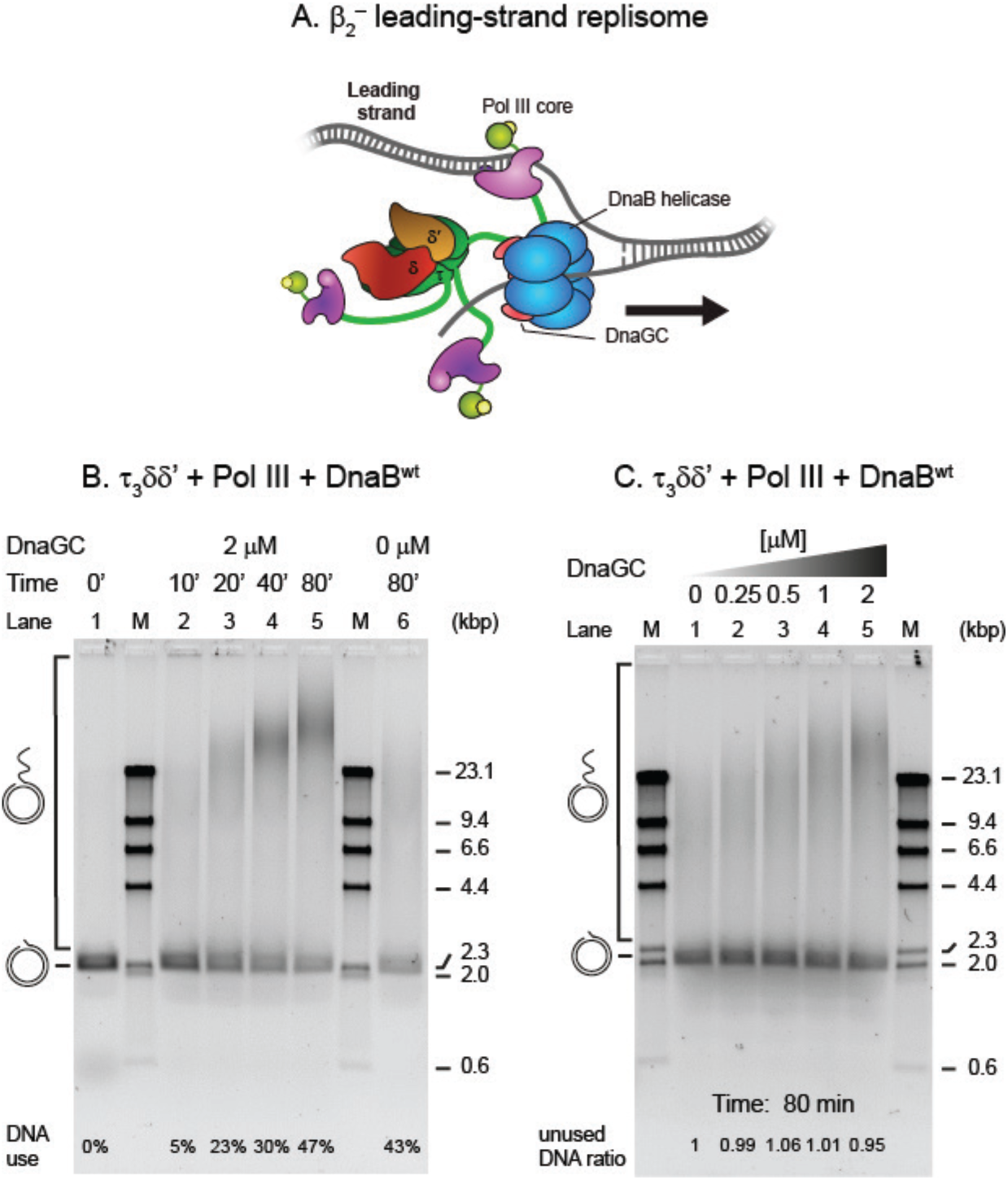
A DnaGC-Induced CLC–DnaB Interaction Stimulates Leading-Strand Synthesis in Absence of the Clamp. (A) Cartoon of the replisome that synthesizes the leading strand only in the absence of the β_2_ clamp, as used in (B) and (C). (B) A time course of rolling-circle DNA synthesis reactions by the β2^−^ replisome, at indicated time points, in the presence of 2 μM DnaGC (lanes 1–5) and in its absence (lane 6) are resolved on a 0.66% agarose gel. (C) Serially diluted DnaGC samples (0.25–2 μM, including zero) were supplemented into the individual rolling circle replication reactions and the replication products visualized on a gel after 80 min (lanes 1–5).

### Binding of Primase Does Not Inhibit DnaB Helicase Activity

It is unknown whether the helicase stalls during priming in *E. coli.* It is also not known how long the primase is bound to the helicase during each OF cycle. As such, it is of importance to know whether the helicase–primase interaction stalls the helicase. Major experimental challenges in characterizing the DNA-unwinding activity of an *E. coli* helicase–primase complex include the multivalency and transience of the interactions and the different helicase conformations. We used a disulfide cross-linked construct, DnaB_6_^F102C^~DnaGC_3_^R568C/C492L^ (DnaB~GC), to remove the stoichiometric heterogeneity of DnaB•(DnaGC)*_n_* (*n* = 0–3) species and to ensure that helicase remains continuously in the dilated state with strong affinity for the CLC. DnaB^F102C^ is a mutant version of DnaB that interacts >200-fold less efficiently with DnaGC. Likewise, DnaGC^R568C/C492L^ interacts with DnaB^wt^ 30-fold more weakly compared to wild type. Nevertheless, the disulfide crosslinking efficiency of these mutants under very mild conditions is nearly 100%, ensuring that for each hexamer of DnaB, essentially all three primase sites are occupied by DnaGC (confirmed by SDS-PAGE and native ESI-mass spectrometry; Lo et al., publication in preparation).

To characterize DnaB~GC, we used SPR and monitored binding of DnaB~GC to immobilized τ_3_CLC. This experiment results in the characteristically slow dissociation (Figure 5A) that we previously observed with DnaB^dilated^ (Figure 2D) or DnaB^wt^ in the presence of DnaGC (Figure 3B). Moreover, similarly to DnaB^wt^ and DnaB^constr^, DnaB^F102C^ alone exhibited a much lower response at equilibrium, accompanied by fast on and off kinetics (Figure 5A). These data indicate that DnaB~GC is indeed mostly in a dilated state and interacts strongly with the CLC.

**Figure 5.**
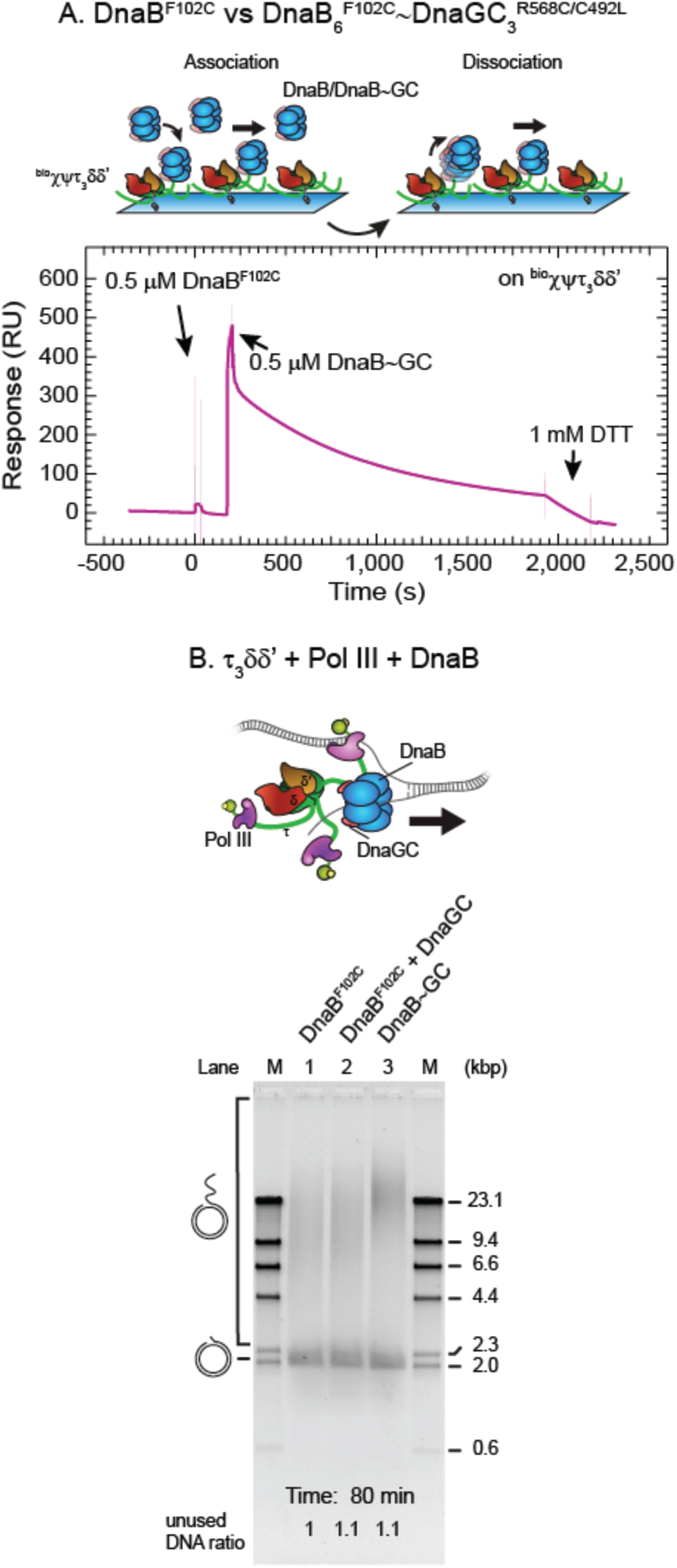
A Cross-Linked DnaB~GC Complex is an Active Helicase. (A) *Top panel*: Cartoon representation of association and dissociation phases of an SPR experiment designed to visualize the binding of cross-linked DnaB~GC to immobilized τ_3_CLC. *Bottom panel*: SPR sensorgrams showing association and dissociation profiles of consecutive injections of DnaB^F102C^ alone and cross-linked DnaB_6_^F102C^~DnaGC_3_^R568C/C492L^ (DnaB~GC) on ^bio^*χ*ψτ_3_δδ’. During DnaB~GC dissociation, dithiothreitol (DTT) injected at ~1,800 s reduces the disulfide cross-link, leading to release of DnaGC and faster dissociation. Spikes in the signal corresponding to imperfect signal subtraction from the control flow cell during solution changes are made more transparent to highlight the relevant portions in the sensorgrams. (B) Rolling-circle leading-strand replication reactions in the absence of β_2_ were supplemented with DnaB^F102^ (lane 1), DnaB^F102^ and 2 μM DnaGC (lane 2), and DnaB~GC (lane 3) and the products were separated on a 0.66% agarose gel following 80 min reaction. See also Figure S5.

We next used DnaB~GC in our leading-strand assay lacking the β_2_ processivity clamp and compared its activity to a DnaB^F102C^ control (with and without DnaGC; Figure 5B). We reasoned that if the activity of the DnaB~GC driven replisome is found to be similar to or stronger than those of controls, it would indicate that DnaB remains an active helicase even when primases are bound to it. The control lanes show that DnaB^F102C^ is active (Figure 5B, lane 1) while the presence of 2 μM DnaGC makes no difference to yield and product length (Figure 5B, lane 2). No change in activity is expected because of the much weaker affinity of the mutant helicase for the primase. However, the presence of DnaB~GC in the reaction results in higher replication efficiency compared to control lanes (Figure 5B, lane 3 *cf.* lanes 1 and 2). Considering that the template consumption was consistent across reactions, our results show that DnaB is an active helicase when it is bound to all three DnaGCs, further confirming that the dilated state is functional. Interestingly, leading-strand assays in the presence of the β_2_ clamp revealed that replisomes with DnaB~GC are not more efficient than those with DnaB^F102C^ (Figure S5A). This result suggests that if Pol III on the leading strand is stabilized by interactions with β_2_, the strengthening of the DnaB–CLC interaction becomes less critical for the stability of β_2_–Pol III^lead^–CLC–DnaB. Taken together, we conclude that DnaB shows helicase activity in both constricted and dilated states (Figure S5B), even when associated with its principal binding partners within the replisome.

### DnaG Concentration Controls the Number of Pol III^∗^s Associated with the Replisome

The replication assays discussed above enabled us to detect functional differences and similarities in partial replisomes as a function of the strength of DnaB–CLC interaction. Next, we set out to examine the influence of the conformational switch in the helicase on full replisome activity by realtime, single-molecule observations of replication. We previously reported the use of a singlemolecule rolling-circle assay to image fluorescently labeled Pol III^∗^ complexes associated with the replisome and visualize dynamic exchange of replisome-bound Pol NI^∗^s with those in solution on the time scale of seconds (Lewis et al., 2017). We hypothesized that [DnaG]-induced strengthening of the DnaB–CLC interaction could contribute to the accumulation of Pol NI^∗^s at the replication fork by slowing down their exchange, and conversely, a weakening could promote exchange and lower the steady-state population.

We employed single-molecule FRAP (fluorescence recovery after photobleaching) experiments (Lewis et al., 2017) and measured the recovery time of fluorescence signal upon photobleaching due to the turnover of fluorescently labeled Pol III^∗^ at the replication fork at different DnaG concentrations (30–300 nM) as both leading and lagging strands are replicated (Figure 6A). The population of labeled Pol III^∗^ in the field of view was photobleached every 20 s with 2-s pulses at high laser power (Figure 6B, top panel) and the recovery of intensity of the fluorescent Pol III^∗^ signal at the fork tracked as a function of time after each high-power pulse (Figure 6B, bottom panel). Fluorescence intensities were converted to numbers of Pol III^∗^s by calibrating the intensity of a single labeled Pol III^∗^, as described before (Lewis et al., 2017). To analyze the data, we pooled together every recovery interval in which we observed replication (Figure 6C; red circles) for a particular DnaG concentration and fit (Figure 6C; blue curve) their averaged fluorescence intensity values (Figure 6C; black squares) with a FRAP recovery equation (Equation 3, Methods). This procedure was repeated for each [DnaG] (Figure 6D and Figure S6A–C).

**Figure 6.**
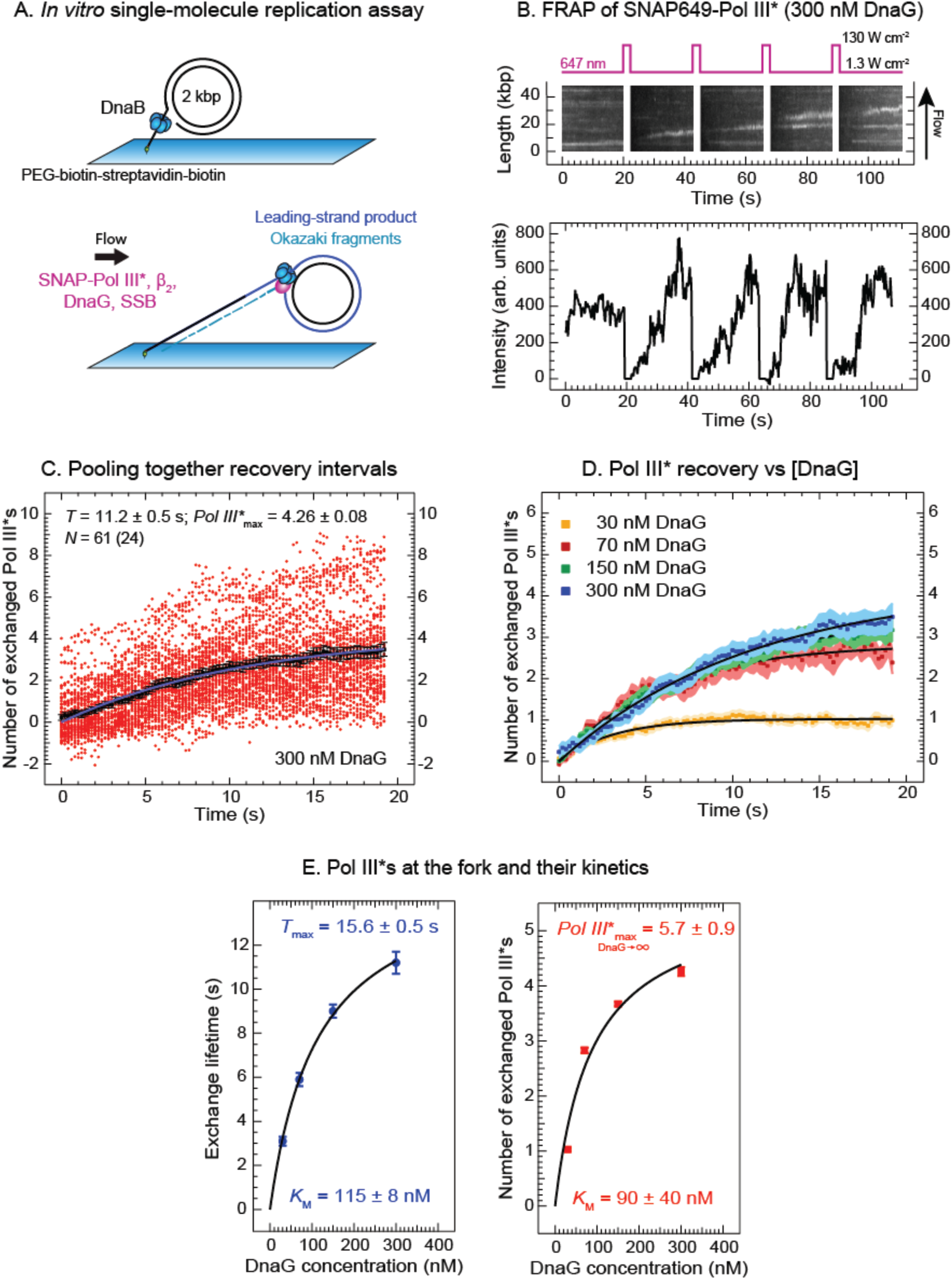
Single-Molecule FRAP Experiments: DnaG Stimulates Accumulation of Polymerases and Slows Their Exchange Dynamics at the Replication Fork. (A) Cartoon showing the two stages of the single-molecule leading- and lagging-strand rolling-circle DNA synthesis assay. First, the 2-kbp rolling-circle substrate with DnaB is loaded with its 5’-tail bound to a coverslip surface through a biotin-streptavidin bond. Then SNAP-Pol III^∗^, β_2_, DnaG, and SSB are introduced to initiate replication. (B) *Top panel*: a representative kymograph of SNAP-Pol III^∗^ at the replication fork in the presence of 300 nM DnaG. Every 20 s, a 2-s high-power pulse laser is used to photobleach the population of SNAP-Pol III^∗^ in the field of view. *Bottom panel*: recovery of SNAP-Pol III^∗^ intensities over time for an individual active replisome. (C) Intensities (red circles) obtained from 61 recovery-interval trajectories of 24 replisomes at 300 nM DnaG are converted into the number of exchanged Pol III^∗^ and displayed, together with their average values (black square). Fitting the evolution of average recovery intensities in time with the FRAP recovery equation (Equation 3, Methods) provides the characteristic (exchange) time (*T* = 11.2 ± 0.5 s), the maximum number of exchanged Pol III^∗^ (*Pol III*^∗^_max_ = 4.26 ± 0.08), and the remaining background intensity *y*_0_, converted in number of Pol III^∗^s (*y*_0_ = 0.72 ± 0.04), then subtracted from every other curve. (D) Averaged recovery intensities at each DnaG concentration (30, 70, 150, and 300 nM) and their FRAP recovery fit curves (black) are shown. The remaining values for the *T*, *Pol III*^∗^_max_, and *y*_0_ are presented in Figure S6. (E) *T* (*left panel*) and *Pol III*^∗^_max_ (*right panel*), plotted as a function of DnaG concentration, are fitted with a steady-state equation (Equation 4, Methods), providing the *K*_M_S (115 ± 8 nM and 90 ± 40 nM, respectively) and either the maximum exchange lifetime (*T*_max_ = 15.6 ± 0.5 s, *left panel*) or the maximum number of exchanged Pol III^∗^ (*Pol III*^∗^_max, DnaG→∞_ = 5.7 ± 0.9, *right panel*) as DnaG approaches infinity.

Our single-molecule analysis revealed that the characteristic exchange time (exchange lifetime, T) is increasing with DnaG concentrations in a concentration range that is physiologically relevant (Figure 6E, left panel). This observation of Pol III^∗^ stabilization at the fork with increasing [DnaG] can readily be explained by DnaB spending more time in a dilated-like state as DnaG levels increase, leading to progressively slower exchange kinetics. In addition, extrapolation of *T* to 0 as [DnaG] approaches zero implies that the translocating DnaB resumes the constricted-like state on DNA as it unwinds dsDNA in the absence of DnaG–DnaB contacts. Further, our data reveal that the number of replisome-associated Pol III^∗^s at 100 nM DnaG, the concentration found in the cell (Rowen and Kornberg, 1978), is two to three, increasing to four copies at 300 nM DnaG (Figure 6E, right panel). Fitting the number of exchanged Pol III^∗^ against [DnaG] to a steady-state equation (Equation 4, Methods) shows that the maximum number of associated Pol III^∗^ can be as high as six (Figure 6E, right panel). The *K*_M_ extracted from these data is 90 ± 30 nM, ~30-fold below the *K*_D_ of the DnaB–DnaG interaction (2.8 μM; Oakley et al., 2005) but reasonably close to the *K*_M_ value for primer utilization (17 ± 3 nM; Graham et al., 2017). This observation suggests that the transition to the dilated-like state depends on additional interactions of DnaG in the replisome, presumably with the lagging-strand template (*i.e.*, during priming events).

## DISCUSSION

In this report, we present evidence for at least two functional modes of interaction between the bacterial clamp loader and helicase CLC–DnaB, corresponding to different conformations of the N-terminal collar in DnaB. We first isolated and characterized CLCs on the surface of an SPR chip (Figures S1 and S2), then demonstrated that binding between the CLC and DnaB^wt^ is weak and transient (Figure 2B and Figure S3C). In solution, DnaB appears to be predominantly in a constricted-like state (Figure 1B, right panel), considering that its affinity for the CLC (Figure 2B) is not much different (~2.5-fold less) from a constricted DnaB mutant (DnaB^constr^) (Figure 2C). In the constricted conformation, one of the two contact regions in the pair of adjacent monomers of the DnaB hexamer responsible for the asymmetric interaction with DnaG is sterically inaccessible (Strycharska et al., 2013). Consistent with this structural picture, the conformationally-constrained DnaB^constr^ did not support priming and OF synthesis (Figure 2F). However, in the dilated conformation, DnaB^dilated^ interacts strongly with the CLC and the complex is slow to dissociate (Figure 2D). The binding of DnaGC promotes a DnaB conformation with high affinity for the CLC, so that binding strength increases >400-fold (Figure 3A and Figure S4E). Nevertheless, the dissociation of DnaB from the CLC is still fast in the absence of DnaGC in solution, whereas DnaB^∗^DnaGC binding was unaffected by the presence of the CLC at high DnaGC concentrations (Figure S4D and Oakley et al., 2005). Given the multivalency of the DnaB•DnaGC interaction, we posit that this outcome is possible only via the positively-cooperative binding of multiple DnaGCs, with the conformational switch initiated by binding of the first DnaGC to DnaB. However, in the presence of high [DnaGC] (5 μM), the dissociation of DnaB from CLC is slow (Figure 3B), similar to the dissociation of DnaB^dilated^ (Figure 2D). We thus conclude that binding of the primase induces the conformational change in the replicative helicase from a constricted-like state with low affinity for the CLC to a dilated-like state with a high affinity for the CLC.

Next, we demonstrated the functional significance of high DnaB•CLC affinity in a modified leading-strand DNA synthesis assay in the absence of the processivity factor β_2_ sliding clamp (Figure 4A). Sliding clamps are utilized in all domains of life to stabilize polymerases as they translocate on DNA between successive nucleotide incorporation steps (Lewis et al., 2016). We hypothesized that the intentional weakening of the Pol III core binding to DNA due to the absence of the clamp might expose the relative change in strength of other contributing interactions, *i.e.* that of DnaB for the CLC. Indeed, this was the case: strengthening the DnaB•CLC interaction in the presence of increasing DnaGC concentrations led to more efficient DNA synthesis (Figure 4B,C), whereas template utilization was not affected (Figure 4C).

The precise mechanism by which the *E. coli* replisome coordinates repetitive lagging-strand priming with DNA synthesis on both leading and lagging strands is not known (Dixon, 2009). One possibility is that the helicase pauses during primer synthesis – a hypothesis based on initial single-molecule studies of the phage T7 replication system (Lee et al., 2006). Another scenario is that the helicase continues unimpeded unwinding of dsDNA at the fork as it remains in contact with the primase during primer synthesis on the lagging strand. This latter mechanism would result in the temporary formation of a “priming loop” that collapses as the new primer is handed off from primase to polymerase (Manosas et al., 2009; Pandey et al., 2009). A critical difference between these mechanisms is the ability of the primase to modulate the helicase activity, in particular whether DnaB helicase pauses when it is bound to primase. To answer this question, we developed a cross-linked DnaB~GC construct (Lo et al., unpublished) and used SPR measurements to show that the binding of DnaB~GC to the CLC is strong and that the dissociation from the CLC is slow (Figure 5A). These observations suggest that the crosslinked DnaB is in the dilated conformation, as expected for the DnaG-bound form. We then tested the activity of DnaB~GC in leading-strand synthesis, both in the absence (Figure 5B) and presence of the clamp (Figure S5A), and confirmed that DnaB remains an active helicase, not only in the constricted conformation (Figure 2F, lanes 5,6), but also in the dilated form that is linked to DnaGC and strongly associated with the CLC (Figure S5B). Our findings thus suggest that the formation of priming loops in *E. coli* is possible, and if it does not happen, there must be a rather specific mechanism in place to prevent it.

Finally, we demonstrated the significance of the DnaG-induced conformational switch in the helicase in the context of full replisomes during active DNA synthesis. We measured the recovery of fluorescence intensity at the replication fork following photobleaching of fluorescent Pol III^∗^ (Figure 6B). Our measurements revealed that the maximum number of replisome-associated Pol III^∗^ increases as DnaG concentration increases from 30 to 300 nM. In the physiological range of [DnaG] (50–100 nM), there are on average 2–3 exchangeable Pol III^∗^ complexes residing at the fork (Figure 6C,D and Figure S6). Fitting the maximum number of associated Pol III^∗^ against DnaG concentration revealed that the upper limit of associated Pol III^∗^ at the fork could be as high as ~6 (Figure 6E, right panel). We cannot exclude that the association of additional Pol III^∗^ bound to the replisome at high polymerase concentrations is caused by weak interactions between Pol III^∗^ and other components of the replisome. Next, a similarity in trends of both the exchange lifetime *T* (Figure 6E, left panel) and the number of associated Pol III^∗^s at the fork (right panel) to increase with rising DnaG concentration can be rationalized by slowing down of CLC•DnaB dissociation when DnaG is bound (Figure 3B *cf.* Figure 2B). These trends suggest that higher DnaG levels increase the portion of time that the helicase spends in dilated-rather than constricted-like states. Conversely, the trend in the fit of *T versus* [DnaG] towards lower concentrations suggests a zero exchange time in the absence of DnaG, implying that under those conditions, the actively translocating helicase resumes a constricted-like state. Using a very different structural and biochemical approach, a model proposed by Strycharska et al. (2013) also predicted that DnaB would be in the constricted state except when it contacts DnaG for priming.

While the single-molecule photobleaching experiments are at physiologically relevant DnaG concentrations, the amount of Pol III^∗^ in those assays (3 nM) is lower than is found inside the cell (~25 nM) (Lewis et al., 2017)]. We chose this lower concentration to enable experimental access to exchange times across the entire range of [DnaG] tested (30–300 nM). The *T* we measured at 300 nM DnaG of 11.2 ± 0.5 s is consistent with previous measurements (11.0 ± 0.6 s) under the same conditions (Lewis et al., 2017). Those measurements also showed that *T* is reduced ~6-fold upon increasing Pol III^∗^ concentration from 3 to 13 nM. At 70 nM DnaG, a concentration approximate to that found in the cell (Rowen and Kornberg, 1978), we measured *T* to be ~6 s; thus, a reduction of *T* to around one second and an increase in the number of exchangeable Pol III^∗^s to >3 would be expected at physiological Pol III^∗^ and DnaG concentrations. These numbers would suggest that the exchange lifetime per individual associated Pol III^∗^ molecule (*T* divided by Pol III^∗^_max_) is well below one second and on par with the cycle time of Okazaki fragment synthesis.

The *K*_D_ value measured for the DnaG–DnaB interaction (2.8 μM) (Oakley et al., 2005) is much higher than the *K*_M_ (90 ± 40 nM) we observe for the DnaG concentration-dependent exchange process. This discrepancy exposes the role of other interactions that DnaG establishes in the replisome that are important for exchange. Taking into account the uncertainty and the different origin of the observables we measure, the fitted *K*_M_ is reasonably close to the *K*_M_ value for primer utilization (~20 nM) measured by others (Graham et al., 2017). These observations underscore the critical role of interaction between DnaG and the lagging-strand template for the primase-induced conformational switch in the helicase. This global CLC•DnaB interaction framework ensures that Pol III^∗^ is not sequestered by helicase in solution and would instead be preferentially bound by a translocating helicase that is engaged in interactions with primase at the apex of the fork.

So why would there be a need for DnaG priming as a signal that triggers a helicase conformational change, which in turn recruits Pol III^∗^s to the vicinity of the replication fork? One obvious possibility is that a Pol III^∗^ newly recruited to DnaB could participate downstream in the primase-to-polymerase switch (Yuzhakov et al., 1999) and subsequent OF synthesis, operating in conjunction with a different Pol III^∗^ that is already replicating the leading strand. Such a model with multiple Pol III^∗^ complexes acting at the fork would allow for a large number of scenarios describing polymerase behaviour in replication, including Pol III^∗^ being left behind on a nascent OF and the simultaneous synthesis of multiple OFs (Geertsema et al., 2014; Duderstadt et al., 2016).

In enzymology, proteins are known to form symmetrical assemblies that undergo cooperative allosteric transitions, thereby serving as switches that allow one molecule in the cell to affect the fate of another (Alberts et al., 2007). Switches are often evolved to enable a tight, ligand-concentration dependent regulation that cannot otherwise be achieved with a single protein. The cooperative allosteric regulation of a DnaG-induced conformational switch in DnaB appears to be capable of functioning as a temporal switch, selected by evolution to support the timely engagement of new Pol III^∗^s into the DNA synthesis process. For example, the simultaneous binding of primase to the exposed site in constricted DnaB and to DNA (*i.e.*, during primer synthesis) enables titration under physiological conditions of the weak (first) DnaG binding site. Subsequently, the helicase undergoes a cooperative allosteric transition to a dilated state and increases its affinity of the remaining two sites for the primase. This behavior in turn greatly increases the affinity of the helicase for the CLC, resulting in the rapid accumulation of Pol III^∗^s at the replication fork. However, upon separation from the primer, primase rapidly separates from one binding site, reversing the initial allosteric transition towards the constricted state and weakening the other two primase binding sites in the helicase. This switch would result in rapid dissociation of Pol III^∗^s from the helicase and its further handoff to a primed site for OF synthesis (primase-to-polymerase switch).

Further work is necessary to deduce whether the proposed primase-to-polymerase switch via the DnaG-induced conformational change in the helicase represents a mechanism that is regularly utilized in OF synthesis, or whether it serves as a backup mechanism to handle roadblocks and obstructions, *i.e.*, when priming on the leading strand becomes necessary or when there are delays in the recycling of the lagging-strand polymerase. The first possibility would appear to be in strongest agreement with the recent paradigm shift in the field proposing a rather stochastic behaviour of Pol III^∗^ at the replication fork, with new Pol NI^∗^s dynamically exchanging in the replisome while DnaB remains stably associated at the fork (Geertsema and van Oijen, 2013; van Oijen and Dixon, 2015; Yuan et al., 2016; Beattie et al., 2017; Lewis et al., 2017; Graham et al., 2017; Monachino et al., 2017). Interestingly, the T7-phage employs a similar strategy of accumulation of polymerases (gp5) on the helicase fused to the primase (gp4) for prompt primer handoff (Loparo et al., 2011; Geertsema et al., 2014). The observation of similar mechanisms in systems of such different complexity suggests that the accumulation of polymerases at the replication fork for primer handoff could be a conserved mechanism in nature.

## Supplemental Information

Supplemental Information includes six figures.

## Acknowledgments

We are indebted to Karl E. Duderstadt and Christiaan M. Punter for ImageJ plugins, Yao Wang for purified proteins, Lisanne M. Spenkelink for development of and assistance with the singlemolecule FRAP assays, and Harshad Ghodke for fruitful discussions. This work was supported by the Australian Research Council (DP150100956 and DP180100858 to A.M.v.O. and N.E.D. and an Australian Laureate Fellowship, FL140100027 to A.M.v.O.), King Abdullah University of Science and Technology, Saudi Arabia (OSR-2015-CRG4-2644 to N.E.D. and A.M.v.O), Nederlandse Organisatie voor Wetenschappelijk Onderzoek (12CMCE03 to E.M.), and the National Institutes of Health (NIGMS R37-071747 to J.M.B.).

## Author Contributions

Conceptualization, E.M., S.J., N.E.D., A.M.v.O.; Methodology, E.M., S.J., J.S.L., N.E.D., A.M.v.O.; Resources, E.M., S.J., J.S.L., Z.Q.X., A.T.Y.L., V.L.O; Software, E.M.; Validation, Formal Analysis, Writing - Original Draft, E.M., S.J.; Investigation, E.M., S.J., J.S.L.; Supervision, S.J., N.E.D., A.M.v.O.; Writing - Review & Editing, E.M., S.J., J.M.B., N.E.D., A.M.v.O.; Funding Acquisition, J.M.B., N.E.D., A.M.v.O.

## METHODS

### Method Details

#### Reagents

Chemicals: (±)-6-Hydroxy-2,5,7,8-tetramethylchromane-2-carboxylic acid (Trolox; Sigma-Aldrich), glacial acetic acid (Ajax Finechem), ADP (Sigma-Aldrich), agarose (Bioline), ATP (Sigma-Aldrich), biotin-PEG-SVA (Laysan Bio, Inc.), catalase (Sigma-Aldrich), dNTPs (dATP, dCTP, dGTP, dTTP) (Bioline), dithiothreitol (Astral Scientific), EDTA (Ajax Finechem), glucose (Sigma-Aldrich), glucose oxidase (Sigma-Aldrich), HCl (Ajax Finechem), K-glutamate (Sigma-Aldrich), MgCh (Ajax Finechem), Mg(OAc)2 (Sigma-Aldrich), mPEG-SVA (Laysan Bio, Inc.), NaCl (Sigma-Aldrich), Na2EDTA (Ajax Finechem), NaOH (ChemSupply), surfactant P20 (GE Healthcare), PDMS (Ellsworth), rNTPs (ATP, CTP, GTP, UTP) (Bioline), SDS (Sigma-Aldrich), Tris (Astral Scientific and Sigma-Aldrich), Tween20 (Sigma-Aldrich).

Gel Electrophoresis: agarose gel loading dye (6x) alkaline (Boston BioProducts), 6x DNA gel loading dye (ThermoFisher Scientific), 10,000x SybrGold (LifeTechnology), GeneRuler DNA Ladder mix (ThermoFisher Scientific), λ DNA/*Hind*III Marker (ThermoFisher Scientific).

#### Buffers

*ALEM buffer*: 2x agarose gel loading dye (6x) alkaline, 200 mM EDTA; *Alkaline buffer*: 50 mM NaOH, 1 mM EDTA; *Imaging buffer*: 30 mM Tris.HCl, pH 7.6, 12 mM Mg(OAc)2, 50 mM K-glutamate, 0.5 mM EDTA, 0.0025% (*v/v*) Tween 20, 0.5 mg/ml BSA, 1 mM freshly made UV-aged Trolox, 0.45 mg/ml glucose oxidase, 0.024 mg/ml catalase, 0.8% (*w/v*) glucose monohydrate, 10 mM dithiothreitol, 1.25 mM ATP, 0.25 mM each UTP, CTP, and GTP, 50 μM each dATP, dTTP, dCTP, and dGTP; *LES buffer*: 2x DNA gel loading dye, 200 mM EDTA, 2% SDS; *Neutralization buffer*: 1 M Tris.HCl, pH 7.6, 1.5 M NaCl; *Replication buffer A*: 30 mM Tris.HCl, pH 7.6, 12 mM Mg(OAc)2, 50 mM K-glutamate, 0.5 mM EDTA, 0.0025% (v/v) Tween 20; *Replication buffer B.* 30 mM Tris.HCl, pH 7.6, 12 mM Mg(OAc)_2_, 50 mM K-glutamate, 0.5 mM EDTA, 0.0025% (v/v) Tween20, 0.5 mg/ml BSA; *SPR1 buffer*: 50 mM Tris.HCl, pH 7.6, 200 mM NaCl, 10 mM MgCl_2_, 0.25 mM dithiothreitol, 0.005% (*v/v*) surfactant P20; *SPR2 buffer*: 25 mM Tris.HCl, pH 7.6, 50 mM NaCl, 5 mM MgCl_2_, 0.25 mM dithiothreitol, 0.005% (*v/v*) P20; *Tris Acetate EDTA (TAE) buffer*: 40 mM Tris, 20 mM acetic acid, 1 mM EDTA (final pH 8.3).

#### Proteins

*E. coli* replication proteins and protein complexes were purified according to previously published protocols. τ_2_γ_1_δδ’, τ_3_δδ’, *χ*ψτ_3_δδ’ (Tanner et al., 2008), DnaB and DnaC (San Martin et al., 1995), γ_3_δδ’ and DnaBC (Jergic et al., 2013), αεθ and SNAP649-αεθ (Lewis et al., 2017), β_2_ (Oakley et al., 2003), DnaB^constr^ and DnaB^dilated^ (Strycharska et al., 2013), DnaG (Stamford et al., 1992), DnaGC (Loscha et al., 2004), DnaB^F102C^ and DnaB~GC (Lo et al., unpublished), and SSB (Mason et al., 2013). Detailed procedures for production of ^bio^*χ* and ^bio^*χ*ψ are described below. Isolation of ^bio^*χ*ψτ_3_δδ’ and ^bio^*χ*ψτ_2_γ_1_δδ’ (Figure S3A) followed methods for production of non-biotinylated CLCs (Tanner et al., 2008).

#### Plasmid construction

*pSJ1376 (^bio^holC)*: The plasmid pET-*χ* (Xiao et al., 1993) that directs overproduction of full-length *χ* was a kind gift of Dr. Mike O’Donnell. It was used as a template for PCR amplification of the *holC* gene using primer 108 (5’-AAA AAA AAC ATA TGA AAA ACG CGA CGT TCT ACC TTC TGG-3’), designed to incorporate a methionine start codon as part of the *NdeI* site, and primer 109 (5’-TTG AAT TCT TAT TTC CAG GTT GCC GTA TTC AGG-3’), that incorporates an *EcoRI* restriction site just following the TAA stop codon. The PCR product was isolated after digestion with *NdeI* and *EcoRI* and inserted between the same set of restriction sites in plasmid pKO1274 (Jergic et al., 2007). Vector pKO1274, a derivative of pETMSCI (Neylon et al., 2000), allows fusion of the gene in-frame behind a N-terminal biotinylation tag: MAGLNDIFEAQKIEWHEH (Beckett et al., 1999) using an *NdeI* restriction site. The resulting plasmid pSJ1376 places the *^bio^holC* gene under the transcriptional control of the bacteriophage T7 ϕ 10 promoter, which directs ^bio^*χ* protein overproduction on addition of isopropyl-β-D-thiogalactoside (IPTG).

#### Overproduction and purification of ^bio^*χ*

*E. coli* strain BL21(γDE3)/pLysS/pSJ1376 was grown at 37°C in LB medium supplemented with thymine (25 mg/l), ampicillin (100 mg/l), chloramphenicol (30 mg/l) and 25 μM (D)-biotin to *A*600 = 0.8. To induce overproduction of ^bio^*χ*, 0.75 mM IPTG was added to the shaking culture. Cultures were grown for a further 3 h, and then chilled in ice. Cells were harvested by centrifugation (11,000 x *g*; 6 min), frozen in liquid N_2_ and stored at –80°C.

After thawing, cells (4.27 g, from 2 l of culture) were resuspended in 65 ml lysis buffer (50 mM Tris.HCl, pH 7.6, 2 mM dithiothreitol, 1 mM EDTA, 20 mM spermidine). The cells were lysed by being passed twice through a French press (12,000 psi). Cell debris was removed from the lysate by centrifugation (30,000 x *g*; 30 min) to yield the soluble Fraction I. Proteins in Fraction I that were precipitated by addition of solid ammonium sulfate (0.45 g/ml) and stirring for 60 min, were collected by centrifugation (35,000 x *g*; 30 min) and dissolved in 30 ml buffer A (30 mM Tris.HCl, pH 7.6, 1 mM dithiothreitol, 1 mM EDTA) containing 150 mM NaCl. The solution was dialyzed against two changes of 2 l of the same buffer to yield Fraction II.

Fraction II was applied at 1 ml/min onto a column (2.5 x 15 cm) of Toyopearl DEAE-650M resin that had been equilibrated against the buffer A+150 mM NaCl. Fractions containing *χ*^bio^ that did not bind to the resin were pooled and dialyzed against two changes of 2 l of buffer A+30 mM NaCl to yield Fraction III.

The dialysate (Fraction III, 50 ml) was loaded at 1 ml/min onto the same DEAE column that had been equilibrated with buffer A+30 mM NaCl. The column was washed with the same buffer and ^bio^*χ* eluted between 1–3 column volumes in a broad peak. Fractions containing ^bio^*χ* were pooled and dialyzed against two changes of 2 l of buffer A+30 mM to give Fraction IV.

The dialysate (Fraction IV, 120 ml) was loaded at 1 ml/min onto a column (2.5 x 10 cm) of Toyopearl SuperQ that had been equilibrated with buffer A+30 mM NaCl. After washing with 60 ml of buffer A + 30 mM NaCl, ^bio^*χ* eluted in a linear gradient (300 ml) of 30–160 mM NaCl in buffer A, in a single peak at ~70 mM NaCl. Fractions containing ^bio^*χ* were pooled and dialyzed against two changes of 2 l of buffer A+30 mM NaCl to give Fraction V.

Fraction V (35 ml) was loaded at 1 ml/min onto a column (2.5 x 15 cm) of heparin-Sepharose 4B (Wijffels et al., 2004) that had been equilibrated against buffer A+30 mM NaCl. The column was washed with the same buffer and ^bio^*χ* eluted between 1–3 column volumes in a broad peak. This purification step did not contribute to improvement in the purity of the protein. Fractions containing ^bio^*χ* were pooled to give Fraction VI.

Proteins in Fraction VI (80 ml) were then precipitated by addition of solid ammonium sulfate (0.45 g/ml) and stirring for 60 min. Precipitated proteins were collected by centrifugation (35,000 x *g*; 30 min), then dissolved in 5 ml buffer A+100 mM NaCl and finally dialyzed against 2 l of the same buffer to yield Fraction VII (6 ml, containing 8 mg of the pure protein).

The molecular weight (MW) of purified ^bio^*χ* determined by nanoESI-MS in 1% formic acid containing 1 mM p-mercaptoethanol (18,650.8 ± 0.2 Da and 18,779.1 ± 0.1 Da) indicated that the N-terminal methionine had been partially removed (in ~80% of proteins) and that biotinylation had not taken place, so the protein was biotinylated *in vitro*, as follows. One part of biomix buffer (50 mM Tris.HCl, 250 mM bicine, pH 8.3, 50 mM ATP, 50 mM magnesium acetate, 250 mM D-biotin) was mixed with three parts of substrate solution (Fraction VII, 70 μM ^bio^*χ*) in buffer A+100 mM NaCl and one part of water, and biotin ligase added to 0.8 μM in final volume of 6.8 ml. This biotinylation mix was then treated at 30°C for 3 h and then dialyzed in 2 l of buffer buffer A+100 mM NaCl at 6°C for storage, yielding Fraction VIII (7.5 ml, containing ~6 mg of ^bio^*χ* in the presence of some biotin ligase). Aliquots were first tested for stability upon freezing in liquid N_2_ and, once the stability on freezing/thawing was confirmed, stored at –80°C.

The MW of *in vitro* biotinylated ^bio^*χ* determined by nanoESI-MS in 1% formic acid, 1 mM β-mercaptoethanol (18,877.1 ± 0.1 Da and 19,005.6 Da) compares well to the calculated value of 18,878 Da in the absence of initiating Met and 19,009 Da when Met is present.

#### Preparation of the ^bio^*χ*ψ complex

Refolding of ψ in the presence of ^bio^*χ* was performed based on methods described by Tanner et al. (2008) with some modifications. About 6 ml of ^bio^*χ* (~4.8 mg) in buffer A+100 mM NaCl was added to 9 ml refolding buffer B (20 mM Tris.HCl, pH 7.6, 100 mM NaCl, 2 mM dithiothreitol, 0.5 mM EDTA) while stirring, followed by drop-wise addition of 1 ml (~10 mg) of ψ in 6 M urea. Consequently, the final concentration of urea in solution was ~0.4 M, a condition that allows ψ to fold and interact with *χ*. The solution was stirred for 4 h at 4°C and then dialyzed overnight in 2 l buffer C (25 mM Tris.HCl, pH 7.6, 90 mM NaCl, 2 mM dithiothreitol, 0.5 mM EDTA, 10% *v/v* glycerol).

Following extensive dialysis, the solution was clarified by centrifugation (35,000 x *g*; 30 min) and the soluble fraction loaded at 1 ml/min onto a column (2.5 x 7 cm) of Toyopearl DEAE-650M that had been equilibrated in buffer C. Fractions containing ^bio^*χ*ψ that did not bind to the column were pooled (12 ml, containing 4 mg of protein complex) and stored at –80°C.

#### Bulk DNA replication assays

Ensemble-averaged (bulk) DNA replication experiments aimed at comparing activities of DnaB^dilated^ and DnaB^constr^ in the context of both leading-strand synthesis and simultaneous leading- and lagging-strand synthesis were commenced by mixing on ice 3.8 nM biotinylated flap-primed 2-kb circular DNA template (Monachino et al., 2018), 1.25 mM ATP, 250 μM each UTP, CTP, and GTP, 200 μM each dATP, dTTP, dCTP, and dGTP, 30 nM *χ*ψτ_3_δδ’, 90 nM Pol III, 200 nM β_2_, 50 nM SSB, 60 nM DnaB^wt^, DnaB^constr^ or DnaB^dilated^, and 360 nM DnaC in replication buffer A in 10 μl reaction volume. When specified, 300 nM DnaG was used for RNA priming on the lagging strand, allowing lagging-strand synthesis to proceed. Since DnaB^constr^ does not load efficiently in the presence of SSB (not shown), the reactions were initiated in a water bath at 37°C for 1 min in absence of SSB to preload helicase. Following the addition of SSB, reactions were incubated at 37°C for further 14 min. In this way, differences in efficiencies of DNA synthesis among the reactions containing different DnaB versions are mainly due to the elongation phase and not to the loading efficiency. Reactions were quenched by mixing equal volumes of replication solution with ALEM buffer, followed by heating in a water bath at 70°C for 5 min and prompt incubation on ice for at least 3 min. Reaction products were then resolved by alkaline agarose gel electrophoresis in a 0.5% (*w/v*) agarose gel. GeneRuler DNA Ladder mix (4 μl) in 1x agarose gel loading dye (6x) alkaline (final volume: 12 μl) was loaded as markers. The alkaline agarose gels were soaked for at least 1 h in alkaline buffer before the reaction products and the markers were loaded. Gels were run at 15 V for ~15 h in a Mini-Sub Cell GT System (Bio-Rad). Then, gels were neutralized in neutralization buffer for ~2 h, and stained with 1x SybrGold in 2x TAE buffer for 7 h. The SybrGold-stained DNA molecules were detected with a Bio-Rad Gel Doc XR (302 nm trans-UV light; Figure 2F).

Bulk leading-strand replication reactions in the absence of β_2_ clamps were assembled by mixing on ice 3.8 nM biotinylated flap-primed 2-kb circular DNA template (Monachino et al., 2008), 1 mM ATP, 400 μM each dATP, dTTP, dCTP, and dGTP, 30 nM τ_3_δδ’, 90 nM Pol III, 30 nM DnaB^wt^, DnaB^F102C^ or DnaB~GC, and unless otherwise stated, 10 mM dithiothreitol in replication buffer A in 10 μl final reaction volume (Figure 4B,C). DnaGC concentration is declared in each experiment. Dithiothreitol was omitted in the reactions that compare the activities of DnaB~GC with DnaB^F102C^ to avoid reduction of the disulfide crosslink in DnaB~GC (Figure 5B). Unless differently declared, reactions were incubated in a water bath at 30°C for 80 min, then quenched by mixing equal volumes of replication solution and LES buffer. Reaction products were separated by gel electrophoresis in 0.66% (*w/v*) agarose gels. λ/*Hind*III DNA digest was loaded as marker. Gels were run in 2x TAE buffer for 100 min at 75 V in a Mini-Sub Cell GT System (Bio-Rad), followed by staining with 1x SybrGold (LifeTechnology) in 2x TAE buffer for 2 h.

Time course bulk leading-strand synthesis reactions with DnaB^F102C^ and DnaB~GC in the presence of β_2_ were assembled as follows: a 60 μl-mix containing 3.8 nM biotinylated flap-primed 2-kb circular DNA template (Monachino et al., 2018), 1 mM ATP, 400 μM each dATP, dTTP, dCTP, and dGTP, 30 nM τ_3_δδ’, 90 nM Pol III, 200 nM β_2_, and 30 nM DnaB^F102C^ or DnaB~GC was prepared by mixing components in replication buffer A on ice; 10 μl were removed for each condition and mixed with same volume of LES buffer to provide the 0 min time-point. The remaining volumes were transferred in a water bath at 37°C. At indicated time-points, 10 μl were removed for each condition and mixed with the same volume of LES buffer to quench reactions. Reaction products were separated by gel electrophoresis in 0.66% (*w/v*) agarose gels; GeneRuler DNA Ladder mixes were loaded as marker. Gels were run in 2x TAE buffer for 150 min at 60 V in a Wide Mini-Sub Cell GT System (Bio-Rad), followed by staining with 1x SybrGold (LifeTechnology) in 2x TAE buffer for 2 h (Figure S5A).

#### Identification and quantification of DNA bands in gels

Quantification of DNA bands in gels was performed using GE Healthcare Life Sciences “Image Quant TL” (*v*.8.1). Lanes were manually identified. The “rubber band” background subtraction algorithm was used. The bands corresponding to the 2-kb DNA template were manually detected and their intensity was calculated by the software (Figure 4B,C, Figure 5B, and Figure S5A).

#### Surface plasmon resonance (SPR) experiments

SPR experiments were carried out on a BIAcore T100/T200 (GE Healthcare) or on a 6 × 6 multiplex Bio-Rad ProteOn XPR-36 system at 20°C, unless stated otherwise. On the BIAcore, an SA (streptavidin-coated) sensor chip (GE Healthcare) was activated with three sequential injections of 1 M NaCl, 50 mM NaOH (40 s each at 5 μl/min). Likewise, all 36 interaction spots on ProteOn NLC (neutravidin-coated) sensor chip were conditioned with three sequential injections of 1 M NaCl, 50 mM NaOH across six vertical (ligand) flow paths (40 s each at 40 μl/min) and six horizontal (analyte) flow paths (40 s each at 100 μl/min).

The stability tests of CLC on the surface of BIAcore T100 SA-chip were performed at 5 μl/min flow using SPR1 buffer. Measurements were initiated by immobilizing ~700 RU of ^bio^*χ*ψ (133 s injection at 325 s, step 1 in Figure S1A,B), followed by association of τ_3_δδ’ at 100 nM (1,500 s injection at 789 s, step 2 in Figure S1A,C). Dissociation, monitored over the period of more than 5,000 s, was interspersed by injections of: (a) SPR1 buffer + 1 mM ADP (120 s injection at 3,110 s, Figure S1C), (b) 1 M MgCl_2_ (60 s injection at 3,554 s, step 3 in Figure S1A,D), (c) solution of δ (550 nM) and δ’ (900 nM) in SPR1 buffer (75 s injection at 4,010 s, step 4 in Figure S1A,E), (d) 1 M MgCl_2_ (60 s injection at 4,554 s, step 5 in Figure S1A), (e) solution of δ’ (900 nM) in SPR1 buffer (75 s injection at 4,733 s, step 6 in Figure S1A,F), and (f) the same solution as in (e).

Considering that ^bio^*χ*ψ could not efficiently be regenerated on the chip surface, binding kinetics parameters for ^bio^*χ*ψ–*γ*_3_δδ’ and ^bio^*χ*ψ–τ_2_*γ*_1_δδ’ interactions in the absence and presence of ATP (Figure S2) were determined using single-shot measurements of the reaction kinetics on a ProteOn NLC sensor chip, designed to provide detailed kinetic analysis in a single injection cycle. Following immobilization of ^bio^*χ*ψ in the ligand direction, the chip was rotated 90°, and solutions of serially-diluted *γ*_3_δδ’/τ_2_*γ*τ_1_δδ’ samples in each case (0.625–10 nM, including zero) in SPR1 buffer (or SPR1 buffer supplemented with 1 mM ATP) were made to flow simultaneously through all six horizontal (analyte) channels at 30 μl/min for 700 s. Dissociation of bound molecules was monitored in the same buffer over 6,000 s. For each round of measurements, ^bio^*χ*ψ was reimmobilized on a new ligand flow path. The final sensorgrams were unmodified ligand flow path- and zero-subtracted using ProteOn Manager Software (*v.* 3.1.0.6). Equilibrium (dissociation constant *K*_D_) and kinetic parameters (rate constants, *k*_a_ and *k*_d_) were determined by simultaneous fitting of the five sensorgrams per measured interaction from the optimized concentration range using a 1:1 (Langmuir) binding model incorporated in BIAevaluation software (*v*. 4.0.1).

All other interactions that enabled determination of dissociation constant *K*_D_ for CLC–DnaB^wt^ and mutant DnaB versions, in the absence and presence of DnaGC, as well as CLC•DnaB^wt^–DnaGC were performed on a BIAcore T200 instrument. First, ^bio^*χ*ψτ_1_*γ*_2_δδ’ and ^bio^*χ*ψτ_3_δδ’ were immobilized on the two flow cells of a SA sensor chip to 1,400 and 1,450 RU, respectively in SPR2 running buffer supplemented with 0.2 mM ATP for stabilization of CLCs on the sensor chip surface. Binding studies were initiated by sequential injections of solutions of serially-diluted DnaB^wt^ samples from the optimizsed concentration range (0.0625–8 μM, including zero) in buffer SPR2+0.2 mM ADP at 30 μl/min for 30 s. While ADP stabilizes the DnaB hexamer and does not influence measured values of binding at equilibrium (*R*_eq_) compared to ATP, it was found to reduce unspecific interactions at high [DnaB] and formation of species that were slow to dissociate, which becomes a critical contribution because CLCs on the chip surface cannot be regenerated. Sensorgrams were zero-subtracted and *R*_eq_ values, generated by averaging response values in the gray highlighted region from the appropriate DnaB concentration range, fitted against [DnaB] using a 1:1 SSA model incorporated in the BIAevaluation software *v*. 4.01 (Figure 2B,C and Figure S3C):

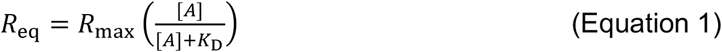

where *R*_max_ corresponds to the response when all the immobilized ligands on the surface are saturated with the analyte A, and [A] is the concentration of analyte in solution.

The τ_3_CLC–DnaB^constr^ interaction was measured under similar conditions, except that the temperature was 25°C (Figure 2C). Binding studies for τ_3_CLC•DnaB–DnaGC interaction were also done under similar conditions by sequential injections of solutions containing fixed (100 nM) DnaB and serially-diluted DnaGC samples from the optimized concentration range (0.0625–4 μM, including zero; Figure S4D) at 30 μl/min for 30 s. Analyses of the data were performed using the SSA model as described above. In contrast, preliminary analysis showed that DnaB^dilated^ binds much more strongly to τ_3_CLC, with much different kinetics parameters, so for illustrative purposes 250 nM DnaB_6_^dilated^ was injected for 400 s under the same conditions used for DnaB^constr^, except that the slow dissociation was recorded for over 2,500 s (Figure 2D).

For the measurement of *K*_DS_ of the ^bio^*χ*ψτ_3_δδ’-DnaB^wt^•DnaGC and ^bio^*χ*ψτ_1_*γ*_2_δδ’-DnaB^wt^•DnaGC interactions, a range of solutions of serially-diluted DnaB^wt^ (0.5–64 nM, including zero) in the presence of 5 μM DnaGC were injected at 30 μl/min for 150 s over immobilized ^bio^*χ*ψτ_3_δδ’ (1,800 RU) and ^bio^*χ*ψτ_1_*γ*_2_δδ’ (1,630 RU), in SPR2 buffer containing 0.2 mM ATP and 0.2 mM EDTA, followed by flow of the same running buffer. Given that we varied [DnaB], whereas the DnaB⁃DnaGC complex and not DnaB titrates the immobilized ligand in the low-nM range of DnaB, and that CLC does not affect the DnaB–DnaGC interaction, the concentration of DnaB⁃DnaGC in solution was obtained by solving the quadratic equation for *x* derived from Equation 2:

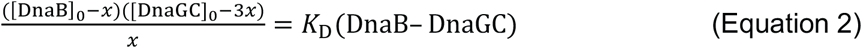

where *x* is solution [DnaB⁃DnaGC], [DnaB]_0_ and [DnaGC]_0_ are the initial concentrations of DnaB and DnaGC (a constant value, equal to 5 μM), and the *K*_D_(DnaB–DnaGC) is the dissociation constant for the interaction of DnaB with DnaGC (1.74 ± 0.09 μM; Figure S4D). This equation is derived based on 1:1 interaction between the DnaB and the first weakly-bound molecule of DnaGC, a property previously also observed by Oakley et al. (2005), in the background of positive cooperative interaction with the second and third DnaGC that each bind > 10-fold more strongly compared to the first one (three DnaGCs allosterically bind to DnaB). Given the appropriate range of 0.5–32 nM of serially-diluted [DnaB]_0_ flowed over τ_3_CLC or 0.5–64 nM in case of τ_1_CLC, Equation 2 was used to calculate the concentration of solutions of DnaB⁃DnaGC samples that were injected over the immobilized ligand: 0, 0.4, 0.7, 1.5, 3.0, 5.9, 11.8 and 23.7 nM in case of τ_3_CLC (Figure 3A), and an extra 47.1 nM injection in case of nCLC (Figure S4E).

To study how the presence of DnaGC affects dissociation of DnaB from CLC, 32 nM DnaB^wt^ with 5 μM DnaGC were injected over immobilized τ_3_CLC in buffer SPR2 containing 0.2 mM ATP and 0.2 mM EDTA at 5 μl/min for 60 s, and then the solution of 5 μM DnaGC in the same buffer coinjected for 1,000 s immediately following the protein association phase to monitor the dissociation of DnaB. Finally, the remaining proteins were washed with the SPR2 running buffer (Figure 3B). In addition, similar conditions were used to test binding of DnaB^F102C^ or disulfide-bond cross-linked DnaB~GC with immobilized τ_3_CLC, except that the proteins (0.5 μM each) were subsequently injected at 30 μl/min for 30 s, and the dissociation of DnaB~GC monitored for 1,720 s, followed by injection for 250 s of the buffer in use that was additionally supplemented with 1 mM dithiothreitol to uncouple DnaB^F102C^ from DnaGC^R568C/C49L^, stimulating the dissociation of DnaB^F102C^ (Figure 5A).

#### *In vitro* single-molecule fluorescence recovery after photobleaching (FRAP) experiments

*In vitro* single-molecule FRAP experiments were performed similarly to a previously published investigation (Lewis et al., 2017). Briefly, a microfluidic flow cell was obtained by positioning a PDMS flow chamber on top of a PEG-biotin-functionalized microscope coverslip. To reduce nonspecific interactions of proteins and DNA molecules with the surface, the chamber was blocked with replication buffer B. The flow-cell was placed on an inverted microscope (Nikon Eclipse Ti-E, Japan) with a CFI Ap TIRF 100x oil-immersion TIRF objective (1.49 NA, Nikon, Japan) and connected to a syringe pump (New Era Pump Systems Inc., Adelab Scientific, Australia) for flow of buffer. Flow-cell temperature was maintained at 31°C by an electrically heated chamber (Okolab, Burlingame, CA).

Leading- and lagging-strand replication experiments were performed under continuous presence of all proteins except DnaBC and as a function of [DnaG] in the 30–300 nM range. Briefly, 40 nM DnaBC were incubated with 340 pM biotinylated flap-primed 2-kb circular DNA template (Monachino et al., 2018), 10 mM dithiothreitol, and 1 mM ATP in replication buffer B for 3 min in a water bath at 37°C. This mixture was then diluted 10 times to a final volume of 220 μl, and loaded into the flow cell first at 40 μl/min for 2.5 min, then at 10 μl/min for 7.5 min, and finally in absence of flow for few minutes (Figure 6A, top panel). During the loading process, the imaging buffer was made. SNAP-Pol III^∗^ was assembled *in situ* by incubating 280 nM τ_3_CLC with 850 nM SNAP649-Pol III in imaging buffer for 90 s in a water bath at 37°C. Finally, the replication solution, which contained 1 mM freshly made UV-aged Trolox, 0.45 mg/ml glucose oxidase, 0.024 mg/ml catalase, 0.8% (*w/v*) glucose monohydrate, 10 mM dithiothreitol, 1.25 mM ATP, 0.25 mM each UTP, CTP, and GTP, 50 μM each dATP, dTTP, dCTP, and dGTP, 3 nM SNAP-Pol III^∗^, 40 nM β_2_, 250 nM SSB, and 30, 70, 150, or 300 nM DnaG in replication buffer B, was loaded into the flow cell first at 20 μl/min for 3.5 min, then at 10 μl/min until the end of the experiment (Figure 6A, bottom panel).

The fluorescently-labeled Pol III^∗^ was visualized by excitation with a 647 nm laser (Coherent, Obis 647–100 CW) at 1.3 W/cm^2^ (photo-bleaching lifetime was 40 s) with an exposure time of 200 ms. Every 20 s, every Pol III^∗^ in the field of view was photo-bleached with a 2-s pulse at 130 W/cm^2^ (photo-bleaching lifetime was 0.7 s) (Figure 6B, top panel). Imaging was done with an EMCCD (Photometrics, Tucson, AZ; 512 Delta). The camera allowed a resolution of 160 nm/px. Because of the high efficiency of *E. coli* DNA replication, only one field of view per experiment was recorded for 5 min. Each experimental condition was performed at least twice.

The analysis was done with Fiji, using in-house built plugins and macros. Briefly, using Fiji, every field of view was corrected to account for background and beam profile. Individual replicating DNA molecules were manually located. Then, the position of the fluorescently-labeled Pol III^∗^ at the tip was semi-manually tracked (Figure 6B, top panel) and its integrated intensity calculated in a 5-px-by-5-px square, applying a local background subtraction (Figure 6B, bottom panel). By calibrating the intensity of a single SNAP649-Pol III, we could convert intensities into number of Pol III^∗^s. At a fixed DnaG concentration, only those recovery intervals where replication occurred were averaged together and fit with the following FRAP recovery function (Equation 3):

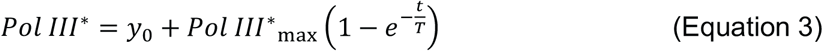

where *Pol III*^∗^_max_ is the maximum number of exchanged *Pol III*^∗^, *T* is the characteristic exchange time, and *y*_0_ accounts for incomplete background removal and was further subtracted from recovery intervals (Figure 6C,D and Figure S6A–C). We averaged recovery intervals from at least 21 individual DNA molecules. Error bars to the averaged values are standard error of the mean (normalized to the number of recovery intervals). The resulting *Pol III*^∗^_max_ and *T*, plotted against DnaG concentration, were fit with a steady-state affinity function (Equation 4):

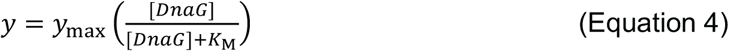

where *y*_max_ represents the maximum value reached by *y* (either *Pol III*^∗^_max_ or *T*) when the concentration of DnaG approaches infinity, while *K*_M_ is a pseudo-Michaelis constant (Figure 6E).

## Supplemental Figures

**Figure S1.**
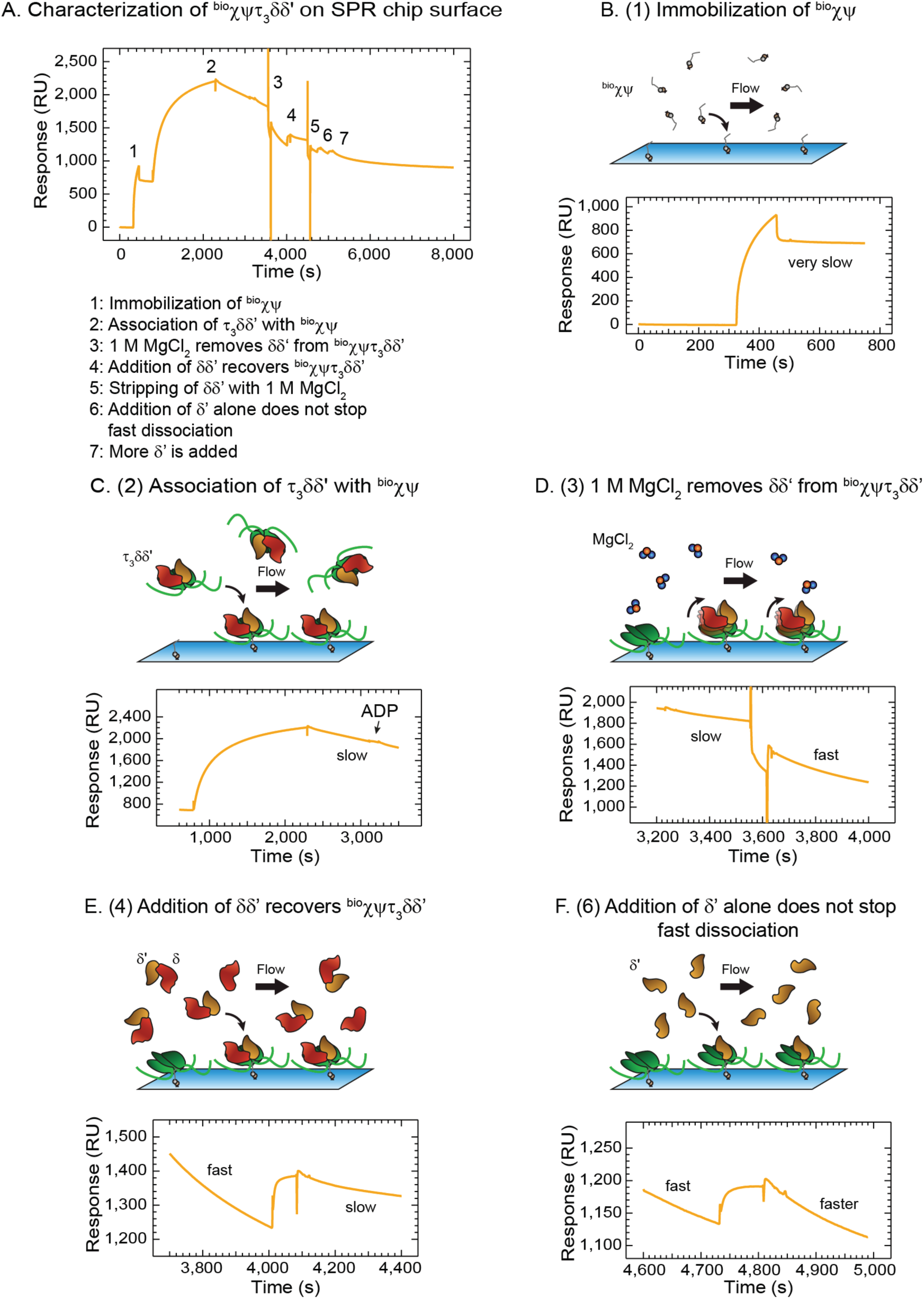
Characterization of the Stability of ^bio^*χ*ψτ_3_δδ’ on SPR chip surface, Related to Table 1. Interrogation of ^bio^*χ*ψτ_3_δδ’ stability by monitoring the association and dissociation phases as various proteins (in SPR1 buffer, containing 200 mM NaCl) or 1 M MgCl_2_ were injected. (A) Steps in the τ_3_CLC assembly and its characterization process. (B) First, ^bio^*χ*ψ was immobilized on the surface. Immobilised ^bio^*χ*ψ proteins are stably bound and dissociation was very slow. (C) To assemble the entire CLC on the chip surface, 100 nM τ_3_δδ in SPR1 buffer was injected for 1,500 s, and uninterrupted dissociation, monitored over ~900 s, allowed estimation of the dissociation half-life *t*_1/2_ (~50 min). Injection of 1 mM ADP did not affect the dissociation rate. (D) Injection of 1 M MgCl_2_ stripped only part of the mass from the surface and accelerated the dissociation rate ~5-fold (*t*_1/2_ ~ 10 min). (E) A 75 s injection of saturating δδ’ assembled *in situ* from 550 nM δ and 900 nM δ’ allowed restoration of a slow dissociation rate (*t*_1/2_ ~ 45 min). (F) In contrast, the injection of 900 nM δ’, which interacts with τ_3_ (Park et al., 2010), following re-treatment with 1 M MgCl_2_ could not restore the slow dissociation rate. These measurements are consistent with the wholesale dissociation of the τ_3_δδ’ complex from immobilized ^bio^*χ*ψ in SPR1 buffer and, surprisingly, with the removal of δδ’ from ^bio^*χ*ψτ_3_δδ’ and dissociation of τ_3_ from ^bio^*χ*ψ following treatment with 1 M MgCl_2_.

**Figure S2.**
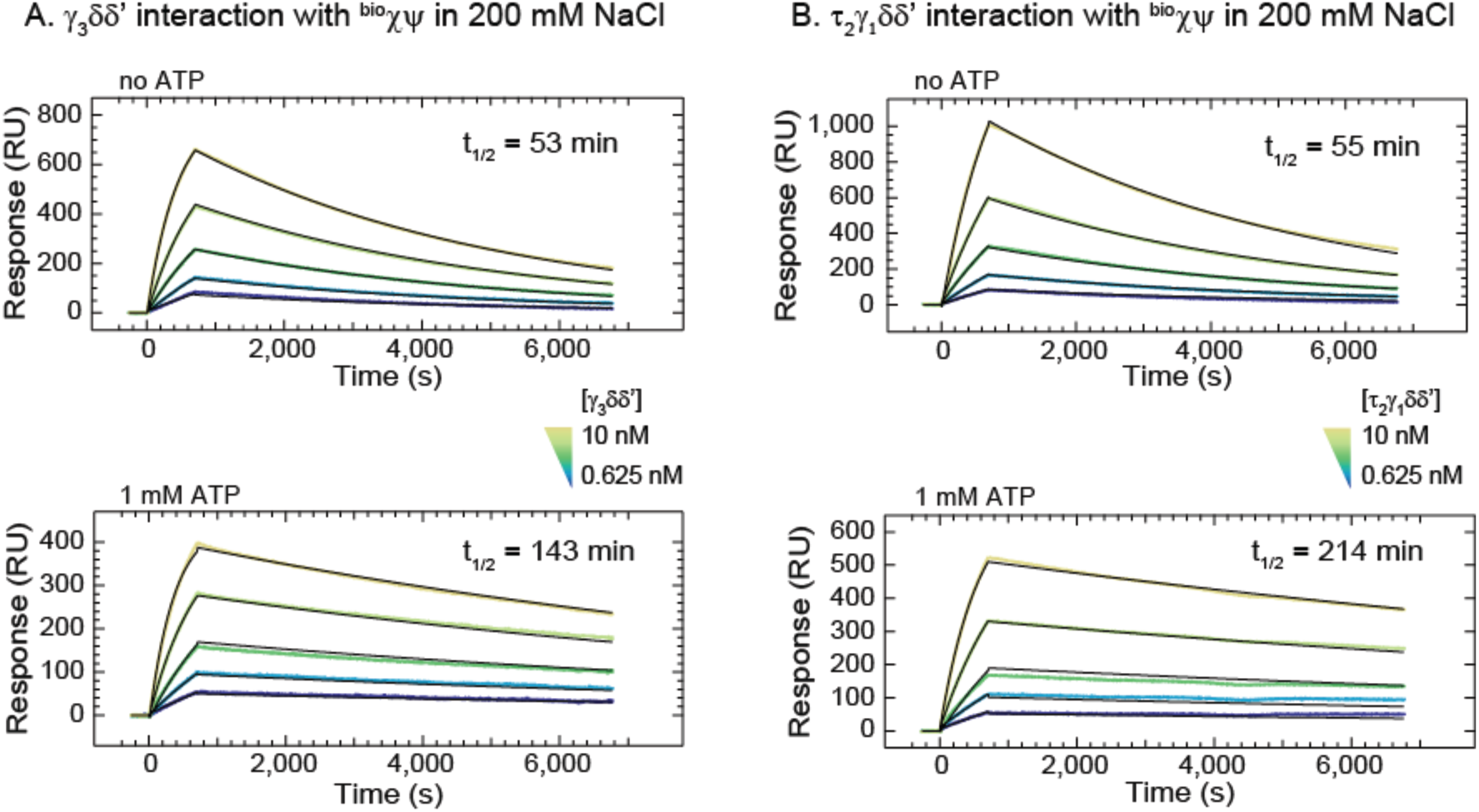
ProteOn Sensorgrams Showing Association and Dissociation Phases of ^bio^*χ*ψ–*γ*_3_δδ’ and ^bio^*χ*ψ–τ_2_*γ*_1_δδ’ Interactions, in the Presence or Absence of ATP, Related to Table 1. (A) Solutions (0.625–10 nM, including zero) of *γ*_3_δδ’, with or without ATP, were injected over immobilized ^bio^*χ*ψ for 700 s, and dissociation was monitored over 6,000 s. Each groups of five sensorgrams (shown in colours) were simultaneously (globally) fit (black curves) using a 1:1 (Langmuir) binding model to determine the corresponding binding parameters *K*_D_, *k*_a_, *k*_d_, and *t*_1/2_ values (Table 1). Errors are standard errors to the fit. (B) The same experiment repeated with the same concentrations of the τ_2_*γ*_1_δδ’ CLC.

**Figure S3.**
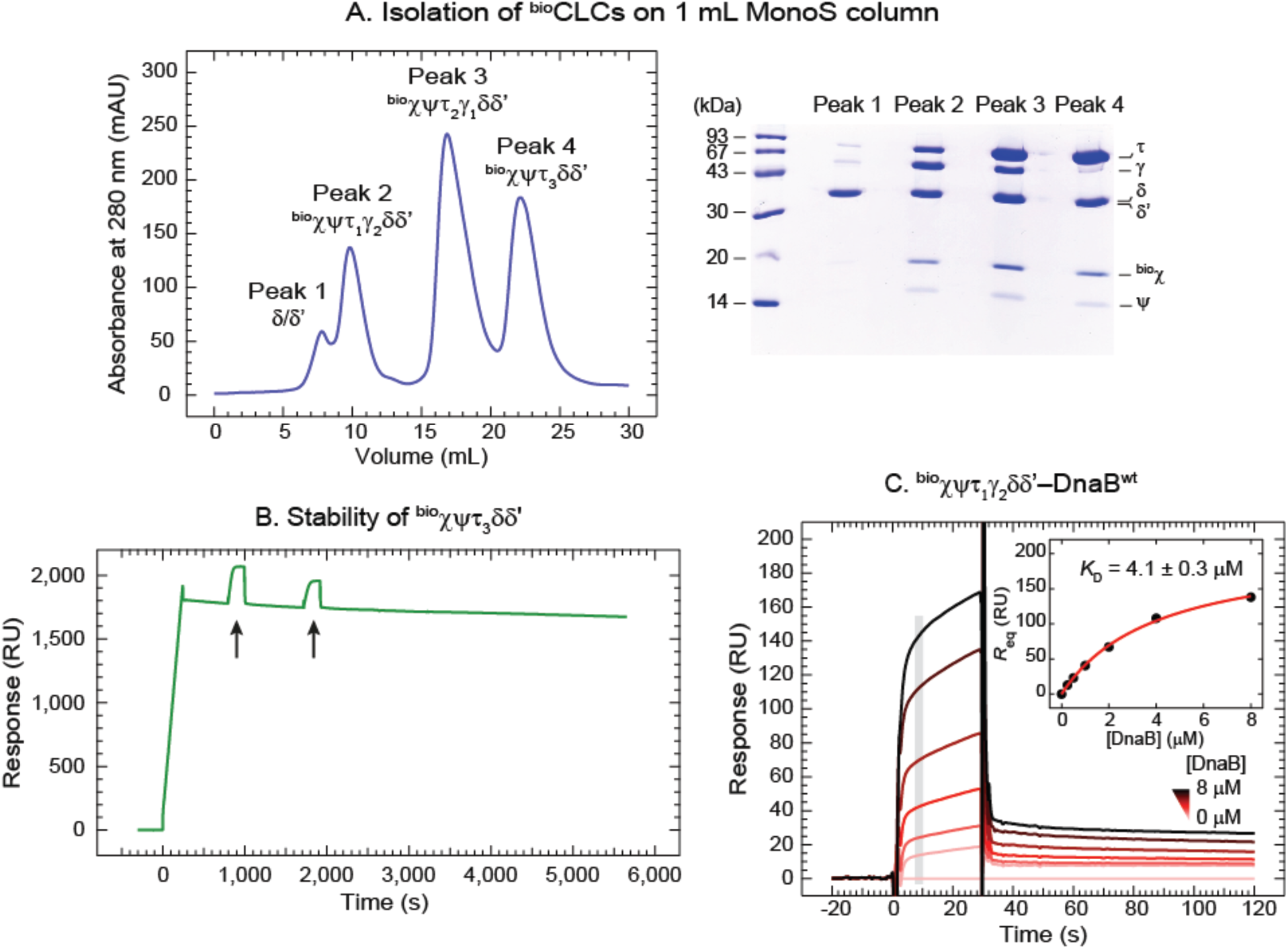
Isolation of ^bio^CLCs and Their Use in SPR Studies, Related to Figure 2. (A) Isolation of ^bio^*χ*ψτ_1_*γ*_2_δδ’, ^bio^*χ*ψτ_2_*γ*_1_δδ’ and ^bio^*χ*ψτ_3_δδ’ clamp loader complexes on a 1-ml MonoS 5/50 GL column (GE Healthcare). Samples from peaks (left panel) were analysed by 4–20 % SDS-PAGE (right panel). (B) The ^bio^*χ*ψτ_3_δδ’ complex was immobilized on a SA chip surface and its improved stability, manifested in slow dissociation in SPR2 buffer that contained reduced NaCl concentration (50 instead of 200 mM) and 1 mM ATP for stabilization of the complex, was monitored for more than 1 h. Control injections of DnaB samples in the presence of DnaGC, which were followed by prompt and complete dissociation response signatures (denoted by arrowheads), confirmed the practical utility of our experimental strategy. (C) SPR sensorgrams show association and dissociation phases for ^bio^*χ*ψτ_1_*γ*_2_δδ’–DnaB^wt^ interaction over a 0.25–8 μM range of DnaB^wt^ (including a zero control). The responses at equilibrium, determined by averaging values in the gray bar region of the sensorgrams, were fit (inset, red curve) with an SSA model to obtain a *K*_D_ value of 4.1 ± 0.3 μM and an *R*_max_ value of 210 ± 7 RU. Errors are standard errors of the fit.

**Figure S4.**
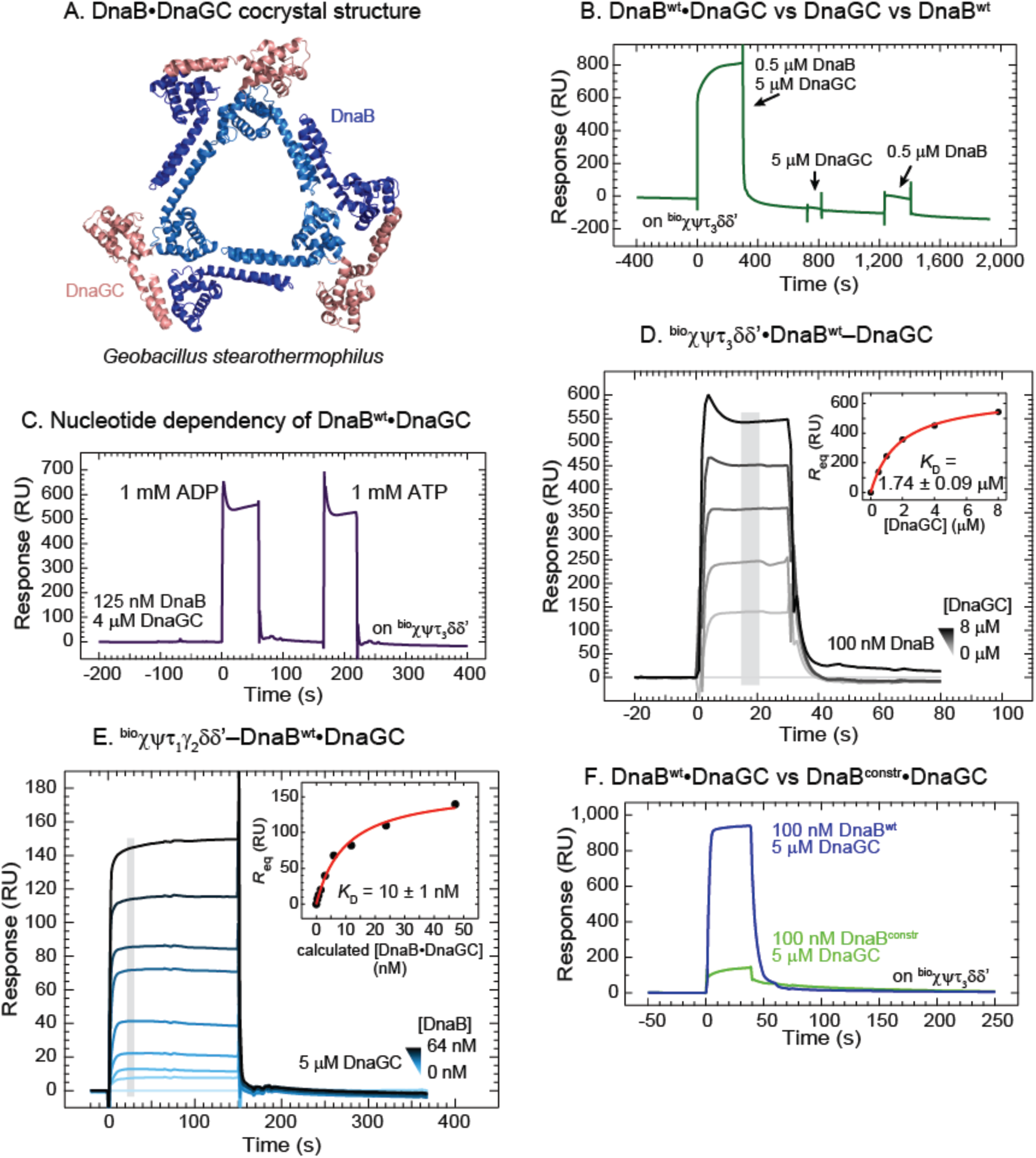
Supplemental SPR Studies of Interactions Between ^bio^CLC and DnaB Variants in the Presence of DnaGC, Related to Figure 3. (A) Co-crystal structure of DnaB_6_•DnaGC_3_ complex from *Geobacillus stearnthermophilus* (Bailey et al., 2007) showing three DnaGC molecules bound to the N-terminal domains of a DnaB hexamer. (B) SPR sensorgram obtained by consecutive injections of 0.5 μM DnaB^wt^ with 5 μM DnaGC, then of 5 μM DnaGC and finally of 0.5 μM DnaB^wt^ in SPR2 buffer with 1 mM ATP shows that while DnaGC stimulates binding of DnaB to immobilized ^bio^*χ*ψτ_3_δδ’, it does not specifically or non-specifically bind to the clamp loader complex at a concentration of 5 μM. (C) Under similar experimental conditions, there was no critical difference in SPR responses if 125 nM DnaB^wt^ and 4 μM DnaGC were injected with either 1 mM ADP or 1 mM ATP. (D) Sensorgrams showing association and dissociation of DnaB^wt^•DnaGC from ^bio^*χ*ψτ_3_δδ’ over a 0.5–8 μM range of serially-diluted DnaGC samples (including a 0 nM control), and at a constant 100 nM concentration of DnaB^wt^. The responses at equilibrium, determined by averaging values in the gray bar region, were fit (inset, red curve) with a SSA model to obtain a *K*_D_ value for the interaction between DnaGC and DnaB^wt^ bound to ^bio^*χ*ψτ_3_δδ’ of 1.74 ± 0.09 μM and an *R*_max_ value of 660 ± 10 RU. Errors are standard errors of the fit. (E) Sensorgrams showing association and dissociation of DnaB•DnaGC from ^bio^*χ*ψτ_1_*γ*_2_δδ’ over a 0.5–64 nM concentration range of serially-diluted DnaB^wt^ samples (including a 0 nM control), and 5 μM DnaGC. The responses at equilibrium, determined by averaging values in the gray bar region, were fit (inset, red curve) against the calculated DnaB^wt^•DnaGC concentrations (0–47.1 nM – see Methods) with SSA model to obtain a *K*_D_ value of 10 ± 1.0 nM and an *R*_max_ value of 163 ± 7 RU. Errors are standard errors of the fit. (F) Comparison of SPR sensorgrams when 100 nM DnaB^wt^ (blue curve) or 100 nM DnaB^constr^ (green curve) were injected in the presence of 5 μM DnaGC over immobilized ^bio^*χ*ψτ_3_δδ’.

**Supplementary Fig. 5.**
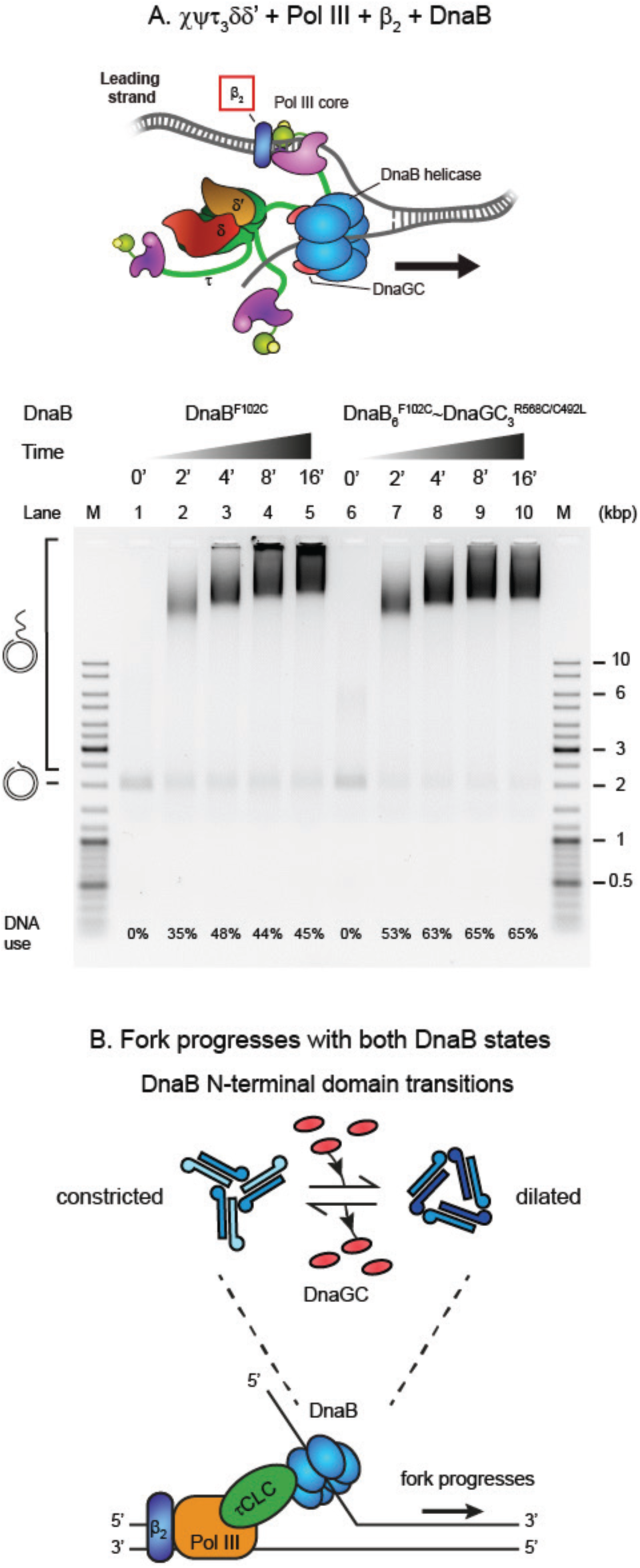
DnaB_6_•DnaGC_3_ Complex and its Effect on Leading-Strand Replication, Related to Figure 5. (A) *Top panel*: cartoon representation of the replisome components required to duplicate the leading strand (leading-strand synthesis). Unlike the replisomes used in the leading-strand DNA synthesis assay shown in Figure 5, the β_2_ clamp is present in the current assay. *Bottom panel*: agarose gel shows DNA products obtained from leading-strand replication as a function of time (0, 2, 4, 8, and 16 min) with either DnaB^F102C^ (lanes 1–5) or cross-linked DnaB_6_^F102C^~DnaGC_3_^R568C/C492L^ (DnaB~DnaGC; lanes 6–10). (B) Cartoon illustration of leading-strand synthesis showing it occurs irrespective of the conformation of DnaB N-terminal domains.

**Figure S6.**
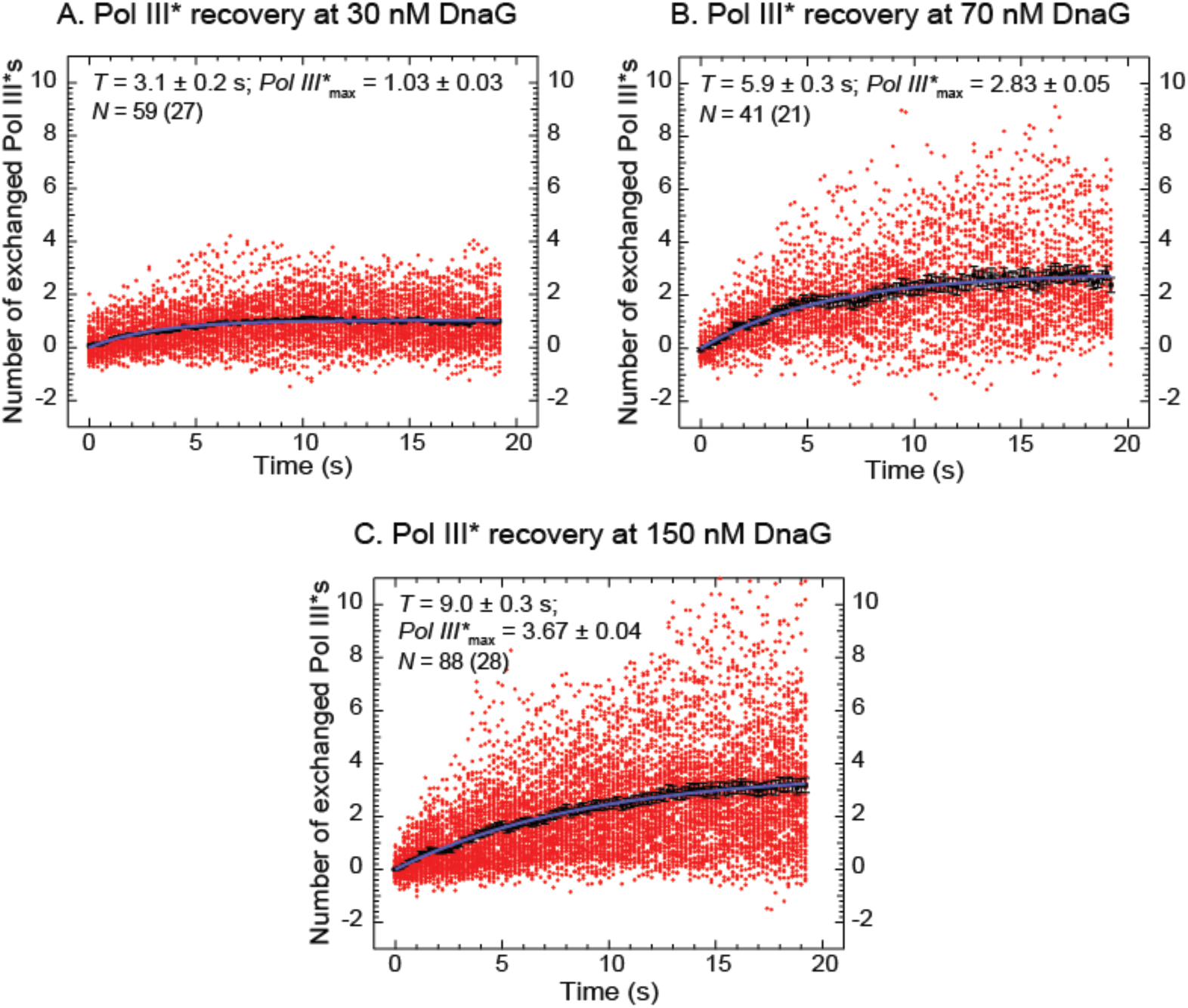
Recoveries of Fluorescence Intensities Following Photobleaching at Different DnaG Concentrations, Related to Figure 6. The intensities (red circles) obtained from the indicated *N* number of recovery-interval trajectories of replisomes (number displayed in brackets) at different concentrations of DnaG (C) are converted into the number of exchanged Pol III^∗^ and displayed, together with their average values (black squares). Fitting the evolution of average recovery intensities in time with the FRAP recovery function (Equation 3, Methods) provided the characteristic (exchange) time (*T*) and the maximum number of exchanged Pol III^∗^ (*Pol III*^∗^_max_), following the subtraction of fit background intensity *y*_0_ that was previously converted into the number of Pol III^∗^. (A) [DnaG] = 30 nM: *T* = 3.1 ± 0.2 s, *Pol III*^∗^_max_ = 1.03 ± 0.03, *y*_0_ = 0.42 ± 0.03. (B) [DnaG] = 70 nM: *T* = 5.9 ± 0.3 s, *Pol III*^∗^_max_ = 2.83 ± 0.05, *y*_0_ = 0.20 ± 0.05. (C) [DnaG] = 150 mM: *T* = 9.0 ± 0.3 s, *Pol III*^∗^_max_ = 3.67 ± 0.04, *y*_0_ = 0.20 ± 0.03.

